# Elastohydrodynamic mechanisms govern beat pattern transitions in eukaryotic flagella

**DOI:** 10.1101/2024.02.04.578806

**Authors:** Shibani Veeraragavan, Farin Yazdan Parast, Reza Nosrati, Prabhakar Ranganathan

**Affiliations:** Department of Mechanical and Aerospace Engineering, Monash University, Clayton, Victoria 3800, Australia; Department of Cell and Molecular Biology, Karolinska Institutet, Sweden

**Keywords:** Flagella, Flagellar beat, Sperm motility, Viscosity sensing

## Abstract

Eukaryotic cilia and flagella exhibit complex beating patterns that change depending on environmental conditions such as fluid viscosity. The mechanism behind these beat pattern transitions remains unclear, although they are thought to arise from changes in the internal forcing provided by the axoneme. We show here that such transitions may arise universally across species via an elastohydrodynamic mechanism. We perform simulations of inextensible and unshearable but twistable Kirchhoff rods driven internally by a travelling bending-moment wave in a fixed plane in the material frame of the rods. We show that, for a large range of beating amplitudes and frequencies, the internally planar driving wave results in the growth of twist perturbations. Outside this domain, the driving leads to simple planar waveforms. Within the non-planar domain, we observe quasiplanar, helical, and complex – perhaps chaotic – beating patterns. The transitions between these states depend quantitatively on physical parameters such as the internal forcing, flagellum stiffness and length, viscosity of the ambient medium, or the presence of a plane wall. Beat pattern transitions in our simulations can be mapped to similar transitions observed in bull and sea urchin sperm when the medium viscosity is varied. Comparison of the simulation results with experimentally observed transitional viscosities in our experiments and elsewhere suggests an assay whereby one can estimate the average force exerted by dynein motors. This could potentially lead to diagnostic assays measuring the health of sperm based on their beating pattern.

**Significance Statement:** The ability of flagella and cilia to manipulate their beating waveforms in different physical environments has important implications for human and animal health. Beat transitions in mammalian sperm flagella, for example, may help sperm navigate the complex reproductive tract to reach the egg. Abnormal beating behaviour, which may have a range of unknown underlying causes, can result in ciliopathies and reproductive disorders. This work provides insight into the physical origins of beat transitions in flagella and shows how they depend quantitatively on flagellum stiffness and length, internal driving provided by the axoneme, viscosity of the ambient medium, and the presence of a plane wall. This could allow one to diagnose the root cause of an abnormal flagellar beat and thus design appropriate treatment.

## Introduction

The rhythmic beating of slender, hair-like appendages in eukaryotic cells is crucial to cellular function. Flagella in free-swimming microorganisms, such as sperm and many algae, play a central role in motility, while cilia in the epithelial linings of larger organisms are essential for sensing and fluid transport (1, 2). Their wavy motion is powered by a highly intricate, conserved internal structure known as the axoneme. While the architecture of the axoneme has been studied in detail, many aspects of the mechanisms underlying its operation remain incompletely understood (3).

In most eukaryotic species, the axoneme consists of a ‘universal’ 9+2 structure: nine interconnected microtubule doublets arranged in a ring-like manner surrounding a central pair of microtubules (3). Dynein motors situated between the doublets in the outer ring exert forces that cause the doublets to slide locally relative to each other. The mechanism that transforms local sliding to global oscillations may also be universal: the seminal work of Julicher and Prost (4) showed that oscillations spontaneously arise as the result of a mechanical instability when a large number of motors act on elastic filaments.

Despite these shared features, swimming eukaryotic cells, particularly sperm, display a remarkable diversity in flagellar beating patterns. For example, bull sperm flagella exhibit transitions from a three-dimensional (3D) rolling beat in bulk fluid to a two-dimensional (2D) ‘slithering’ planar beat when within a micrometre of a surface (5). Similarly, sea urchin sperm exhibit a planar beat close to a wall and quasi-planar beating away from walls (6). Planar beating is also observed when bull sperm swim in fluids of viscosity above around 20 cP (7, 8).

How does one explain these transitions observed in beating patterns? The travelling waveforms generated in the filaments could depend on the biochemical regulation of dynein motor activity, which in turn may modify the mechanism behind the oscillatory instability in the axoneme (9– 17) or modify the amplitude, wavelength, and wave speed of the emergent travelling wave (18– 23). It would therefore seem that the variety in flagellar waveforms is largely due to internal differences in the dynein forcing.

We present simulations that suggest that, like the origin of flagellar oscillations, transitions in beat patterns may also be due to a *universal* elastohydrodynamic mechanism that does not require complex mechanosensing or biochemical regulation of the dynein activity. To do so, we model a bendable and twistable but inextensible and unshearable elastic filament suspended in a viscous fluid. The filament is driven internally by a travelling wave of active bending moments – that is, we take as given a mechanism that generates the internal forcing wave arising from motor activity. Importantly, in every cross-section, the oscillation plane of the forcing wave is kept constant *relative to the material frame of reference associated with that cross-section*. In other words, the forcing wave is *internally planar*. The cross-sections can, however, rotate relative to each other as the filament bends or twists.

We observe that increasing the fluid viscosity can trigger an abrupt transition from planar beating to helical beating and back. We also find that a higher amplitude of the driving wave triggers a transition to helical beating. Our analysis *suggests* that these transitions are due to an elastohydrodynamic instability, indicating that the 2D-to-3D transition in beating patterns can occur *generically* in sperm flagella, and potentially in motile cilia, when the hydrodynamic resistance due to the ambient medium changes, either due to changes in the viscosity of the medium or due to the increased resistance while swimming near a wall. The observations of the transitions in the simulations are in qualitative agreement with the observations of beat transitions in experiments by us (with bull sperm) and others (6–8) (with bull and sea-urchin sperm).

This study thus sheds light on how cilia and flagella could, in general, ‘sense’ and react to the hydrodynamic resistance of their surroundings. These results are relevant to both external and internal fertilization and suggest that the sperm size of different species and other internal operating conditions within flagella may have evolved to ensure a planar beat in their natural environment. Defects in cilia and flagella, such as an abnormal length or bending stiffness, can greatly alter beating patterns and hence lead to poor swimming or impaired function.

## Results

### 3D beating patterns emerge from internally planar oscillations in active bending moments

We modelled a flagellum as an inextensible and unshearable but flexible filament (Kirchhoff rod (24)) suspended in a viscous Newtonian fluid (see Methods) (18, 25). The effect of motor activity was modeled by imposing a sinusoidal travelling wave in the active bending moment. We studied the effect of varying the wavelength, frequency, and amplitude of the imposed active moment wave on the beating motion of the filament. These driving parameters are expressed here in dimensionless terms as the wavenumber, *k*, the frequency *S* = *L*(*ω μ*/*K*_*B*_)^¼^ (which is also known as the Swimming number (26)) and the amplitude ratio, *A* = *αL*/*K*_*B*_, where *L* is the filament length, *K*_*B*_ is its bending stiffness, *¼* is the viscosity of the suspending medium and *α* and *ω* are the forcing amplitude and angular frequency, respectively. The swimming number is the ratio of the time scale of the passive elastohydrodynamic response of the system to *ω*^−1^, the time period of the forcing wave. The order of magnitudes for *S* (from 2 to 21) and *A* (from 5 to 30) were chosen based on the beat frequencies and flagellar curvatures of sperm observed in our experiments. The dimensionless wavenumber *k* was set to either 2π or 4π to result in one or two forcing waves over the length of the filament, respectively. These correspond to the wavenumbers observed in bull and sea urchin (*Echinus esculentus*) sperm. In each case, we studied the distributions of the curvature *κ* and the torsion *τ* along the filament length after 100 forcing oscillations had been imposed. Proper Orthogonal Decomposition (POD) was used to resolve the curvature distribution into spatial modes and their respective time-dependent coefficients (27). The power spectral densities (PSDs) of the first time-dependent POD coefficient of curvature were normalized by their peak values and used to compare the periodicities in the observed beat patterns.

We uncovered four different types of beating patterns (Fig. 1 A-D) by comparing kymographs of the curvature (Fig. 1 E-H) and torsion functions (Fig. 1 I-L), and the curvature PSDs (Fig. 1 M-P). The imposed travelling wave in the active moment resulted in travelling waves emerging in curvature and torsion, as shown by the inclined bands in the kymographs. Since the oscillation in the imposed bending moment is always exerted about the same axis at each cross-section, the simplest beating pattern we expect to emerge in response is *planar* beating, which is characterized by the absence of torsion (Column underneath Fig. 1A; *S* = 7,*A* = 5,*k* = 2*π*; Supplementary Video 1). *However, we observed that torsion can spontaneously emerge in response to driving that is internally unidirectional*. We observed beating patterns that had banded torsional kymographs but with only a single frequency peak in the curvature PSDs (Column underneath Fig. 1B; *S* = 5,*A* = 5,*k* = 2*π*; Supplementary Video 2). When beating is perfectly planar, the orientation of the global beating plane remains fixed in space. In contrast, beat patterns that include a small-amplitude travelling wave in torsion (Fig. 1B, F, J) exhibit a localized twist that travels down the filament. Although the filament remains nearly planar in shape at any instant, the cumulative effect of this torsional wave is a slow, steady rotation of the global beating plane. We therefore refer to these beating patterns as *quasiplanar* states. In contrast, in *helical* beating (Column underneath Fig. 1C; *S* = 12,*A* = 20,*k* = 4*π*; Supplementary Video 3), a helical wave travels from the head towards the tail while the filament rolls. Unlike the quasiplanar state, torsion in this case is no longer localized — the entire filament is twisted at any given instant. The power spectral density (PSD) of the time series of the spatial curvature profile (Fig. 1O) exhibits peaks at the driving frequency and its higher harmonics, reflecting the more complex global deformation. The higher harmonics lead to the interesting feature of small fluttering vibrations superimposed over the more slowly travelling helical waves.

**Fig. 1.**
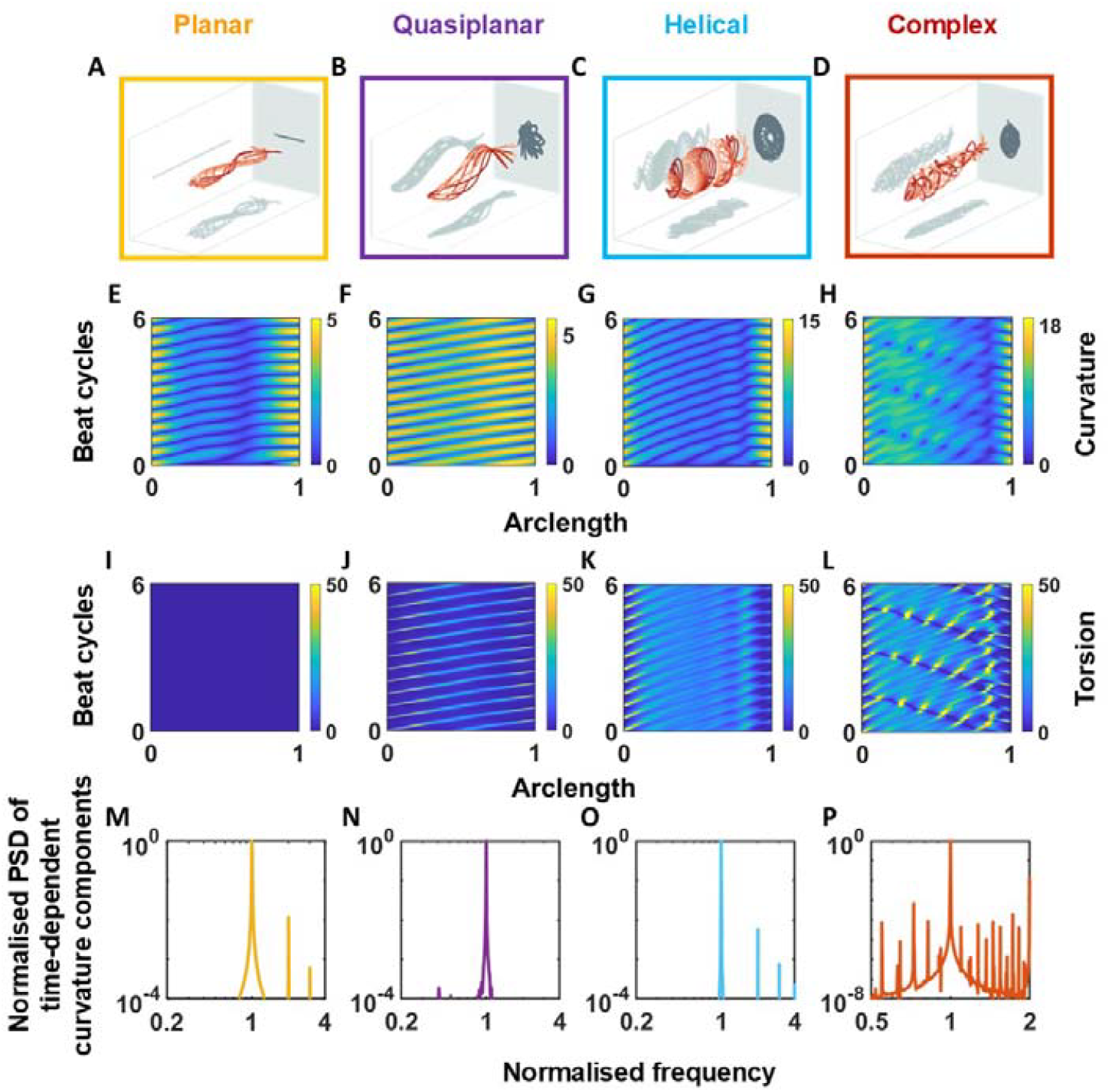
Quantitative comparison of the four beating pattern types. Representative images of the four qualitative beating types: (A) planar (*S* = 7, *A* = 5, *k* = 2π), (B) quasiplanar (*S* = 5, *A* = 5, *k* = 2π),), (C) helical (*S* = 12, *A* = 20, *k* = 4 π), (D) complex (*S* = 15, *A* = 20, *k* = 4 π) beats. (E-H) Curvature kymographs, (I-L) torsion kymographs, and (M-P) normalised power spectral densities of the first time-dependent components of curvature for the representative planar (*S* = 7, *A* = 5, *k* = 2 π), quasiplanar (*S* = 5, *A* = 5, *k* = 2 π), helical (*S* = 12, *A* = 15, *k* = 4 π) and complex (*S* = 15, *A* = 20, *k* = 4 π) beats as labelled. Frequencies are normalized with respect to the driving frequency.

### Can chaotic beating emerge from regular internal forcing?

Beyond planar, quasiplanar, and helical patterns, our simulations revealed a class of beat dynamics that are neither periodic nor simply describable in terms of traveling waves. These *complex* three-dimensional beats (Column underneath Fig. 1D; *S* = 15,*A* = 20,*k* = 4*π*; Supplementary Video 4) exhibit power spectra with both harmonic and subharmonic peaks, and spatiotemporal shapes that fail to repeat from cycle to cycle.

Using simulations based on Resistive Force Theory, we examined the dynamics of a representative complex beat at *S* = 12,*A* = 20 in greater detail and compared them with those of a representative helical beat at *S* = 12,*A* = 15. We restricted our attention to the time dependence of the curvature profile at a single point along the filament (*s* = 0.5). We define the period of a beat cycle to be equal to the period of the forcing oscillation. The dimensionless wavelength in both cases is *k* = 4*π*.

In Fig. 2A, all three curvature components at *s* = 0.5 are plotted against each other over 15 beat cycles (passage of time indicated by color) for the helical beating pattern, beginning after 30 beat cycles from the start of the simulation. In Fig. 2B, we plot the curvatures at *s* = 0.5 at the end of every beat cycle against the curvatures at the end of the subsequent beat cycle to construct a Poincaré-like map of the same beat. This map shows three points — one for each curvature — overlapping over 50 cycles, indicating that the beat is completely periodic.

**Fig. 2.**
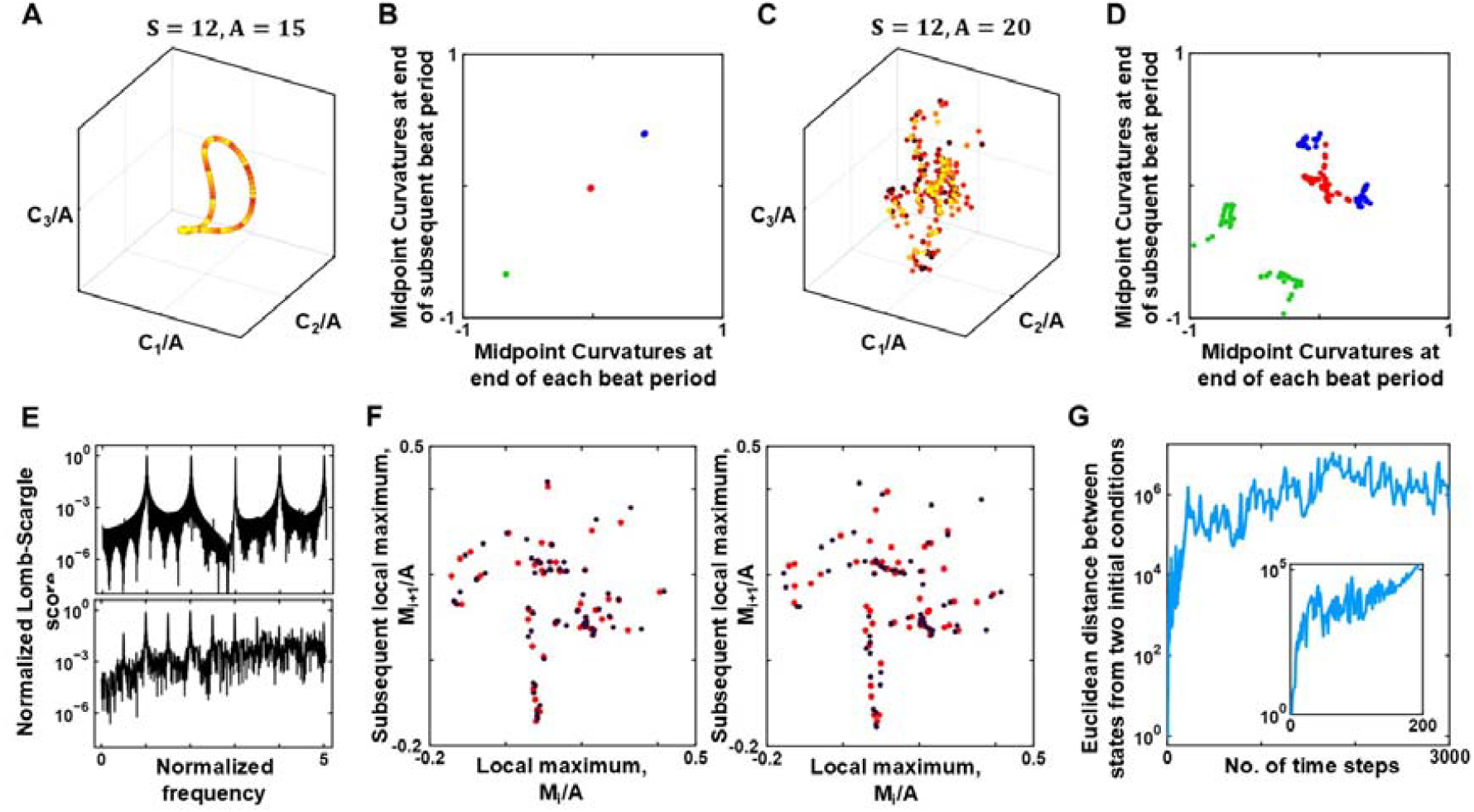
Complex beats are aperiodic and sensitive to initial conditions. **(A)** Curvatures at *s* = 0.5 plotted against each other over 15 beat cycles, beginning after 30 cycles from the start of the simulation for a helical beat at *S* = 12, *A* = 15, *k* = 4π. The passage of time is indicated by color (increasing redness). (B) Curvatures at *s* = 0.5 at the end of every beat cycle plotted against the curvatures at the end of the subsequent beat cycle over 50 beat cycles, for the helical beat parameters in (A). Red points represent C_1_, blue points represent C_2_, and green points represent C_3_. (C) Curvatures at *s* = 0.5 plotted against each other as in (A), for a complex beat at *S* = 12, *A* = 20, *k* = 4π. (D) Curvatures at *s* = 0.5 at the end of every beat cycle plotted against the curvatures at the end of the subsequent beat cycle over 50 beat cycles, for the complex beat parameters in (C). (E) Lomb-Scargle frequency plots for the helical beat (top) and complex beat (bottom) in (A) and (C), respectively. (F) Lorenz maps of the twist, C_1_ at *s* = 0.5 for the two initial conditions (indicated by different colors) between t = 35–60 beat cycles (left) and t = 75–100 beat cycles (right). (G) Time evolution of the Euclidean distance between the states of the system obtained from two different initial conditions, normalized with respect to the initial distance, d ≈ 4.5 × 10^-4^.

Similar plots for the complex beat in Figs. 2C and 2D show scattered points with nearly no overlap, demonstrating that it is not periodic over the time scale of the simulation. The points remain scattered even when only considering the last 15 beat cycles of the simulation, indicating that periodic beating is not achieved even after 80 beat cycles from the start of the simulation.

We used a Lomb-Scargle frequency plot (Fig. 2E) to search for traces of periodicity in the twist profile, considering only the local maxima and minima for simplicity. The helical beat (top plot in Fig. 2E) shows peaks at integer multiples of the driving frequency, as expected. The complex beat (bottom plot) shows additional peaks at non-integer multiples, and numerous small peaks throughout the spectrum. This could indicate quasiperiodicity or chaotic dynamics.

We then repeated the simulation for a nearby initial condition such that the Euclidean distance between the two initial conditions, 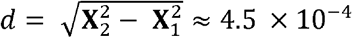, where **X** represents the state of the system at *t* = 0 (see Methods). Fig. 2F shows Lorenz maps of the twist profile at *s* = 0.5 for the two initial conditions (indicated by different colours) between t = 35–60 beat cycles (left) and t = 75–100 beat cycles (right). Comparing the plots, we see visually that the points overlap less in the later beat cycles. Fig. 2G shows the time evolution of the distance *d*, normalized with respect to its initial value. The inset shows approximately linear sections of the plot, which indicate that the distance between the states increases approximately exponentially before saturating due to the finite size of the filament. Together, these analyses indicate that the complex beat *may* exhibit deterministic chaos arising solely from internal mechanical forcing, without external noise or regulation. Such emergent behavior has been previously observed in low-Reynolds-number models of active flexible filaments (28) but has not been systematically examined.

### 2D-3D beat transitions are correlated with changes in medium viscosity

The beat patterns in our simulations resemble those observed in sperm. Planar beating has been observed in various species of mammalian sperm when viscosity is increased (8, 29–31) or close to surfaces (5, 32) as well as in marine invertebrates such as the sea urchin in standard saline (6, 33). The quasiplanar motion is seen in bull (7, 8) and human sperm (30, 31) swimming in low viscosity media (water-like viscosities, Supplementary Video 5) where the flagellum beats in a largely planar fashion and the beating plane rolls as the cell moves. Helical beating in sperm is well known (6), and vibrations in helical beats have also been observed in sea urchin sperm (6) swimming at 1500 cP. Complex non-periodic beat patterns have also been experimentally observed, for example, in pigeon sperm (34).

Since the ratio *S* depends on the viscosity *μ*, our observations (and those elsewhere (35)) suggest a strong connection between beating patterns and medium viscosity. To better understand how viscosity affects beating patterns at any given amplitude of the internal active wave, we mapped the four kinds of beat patterns to the values of *A* and *S* in our simulations for each of the two values of the wavenumbers we studied. When *k* = 2*π* (Fig. 3A), distinct domains are observed corresponding to quasiplanar (purple symbols), planar (yellow), and helical (blue) beats. For values of *S* below 6, the beating pattern was predominantly quasiplanar, accompanied by rolling filament motions. In the region approximately bounded by the lines *S* = 6 and *A* = 1.67*S* − 3, the beat was predominantly helical. At all other values of *S* and *A*, the beat was predominantly planar. The phase map is noticeably different at the higher wavenumber of 4π (Fig. 3B). Specifically, the quasiplanar beat was not observed, and the region of the map over which helical beating occurred diminished to between *S* = 9 and *A* = 2.4*S* − 14.8. At both,*k* = 2*π*, and 4*π*, isolated instances of complex beating occurred at the boundaries between 3D and 2D beats for S and A values higher than 15 and 20, respectively.

**Fig. 3.**
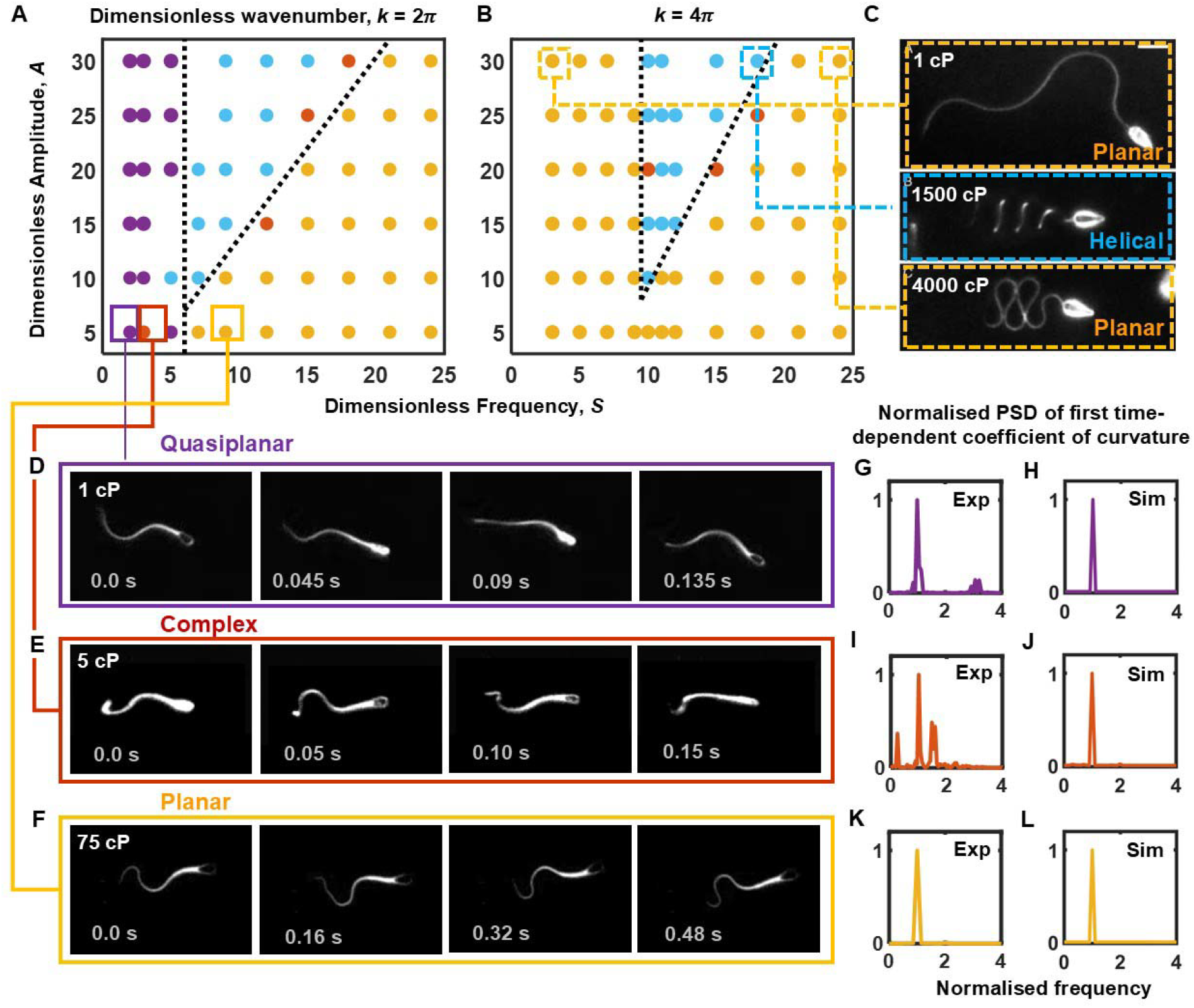
Phase-maps of beating patterns and transitions between them. Phase maps relating the different beating patterns to the amplitude ratio, *A* and the swimming number, *S*, for a wavenumber of (A) 2π and (B) 4π. The yellow, purple, blue and red symbols represent the planar, quasiplanar, helical and complex beat patterns. The dotted boundaries between the phase regimes are drawn to guide the eye. (C) Beat patterns of sea urchin sperm reported by Woolley and Vernon (6) at 1 cP (planar - top), 1500 cP (helical - middle) and 4000 cP (planar - bottom) linked to their equivalent datapoint in A. Time-lapse images of bull sperm from our dark-field microscopy experiments indicating (D) quasiplanar, (E) complex and (F) planar beat observed in external medium viscosity of 1 cP, 5 cP and 75 cP, respectively. Power spectral density (PSD) of the first time-dependent coefficient of the projected 2D curvature from experimental data for (G) quasiplanar beat shown in D, (I) complex beat shown in E, and (K) planar beat shown in F, with the PSDs from our simulations in (H), (J), and (L), respectively.

In the context of sperm flagella, it has been suggested that the bending stiffnesses may be anisotropic due to the 5-6 bridge in the structure of the axoneme (36). While experimental measurements of the bend anisotropy are not available, an estimate of around 2.6 was obtained in a study involving a macroscopic reconstruction of the axoneme (36). We repeated our simulations setting *K*_*B,3*_*/K*_*B,2*_ = 2.6, where *K*_*B,3*_ and *K*_*B,2*_ are the bending stiffnesses along ***d***_**3**_ (along which the internal driving is applied) and ***d***_**2**_ (perpendicular to ***d***_**3**_, on the plane of the filament’s cross section), respectively. The resulting phase map for *k* = 4*π* (shown in Supplementary Figure 1) indicates that the same qualitative beating patterns are observed, although the region over which they occur is diminished. Notably, three-dimensional beating patterns are not observed at *A* = 10 but at *A*= 15 and above. Since our focus is on qualitative behaviour and order-of-magnitude estimates, in the absence of bending anisotropy measurements, we use isotropic bending stiffnesses in the remainder of this paper.

### Elastohydrodynamics explains beat transitions across sperm species

We considered whether our simulation results are consistent with the experimental data of Woolley and Vernon (6) with sea urchin sperm (Fig. 3C), and also our experimental observations with bull sperm (Fig. 3 D-F) swimming (see Methods). In an aqueous medium of 1 cP, where sea urchin sperm exhibit planar beating (Fig. 3C), bull sperm exhibited a rolling quasiplanar beat (Fig. 3D and Supplementary Video 5). The rolling of the cell was observed in the videos as periodic flashes of high intensity in the head region (Supplementary Fig. 2A). The quasiplanarity of the flagellum is also evident in the periodic oscillation of the projected length of the flagellum (Supplementary Fig. 2B): the projected length drops to a minimum (maximum) when the flagellar plane is perpendicular (parallel) to the focal plane. When the medium viscosity was increased to 1500 cP and 4000 cP, sea urchin sperm exhibited helical and non-rolling planar beats, respectively (Fig. 3C). In our experiments with bull sperm, on the other hand, when the viscosity was increased to 5 cP, a complex three-dimensional motion (Fig. 3E and Supplementary Video 6) was observed, whereas at 20 cP and 75 cP, the beat was planar (Fig. 3F and Supplementary Videos 7 and 8).

The flagellar systems studied in experiments are structurally and mechanically more complex than our model: sperm flagella are non-cylindrical and mechanically anisotropic, and sperm cells have heads. However, we propose that our idealized filament model can serve as an *effective* or standard system onto which real biological filaments can be mapped. Rather than attempting to design simulations that match all the anatomical detail, inhomogeneous and anisotropic material properties, and regulatory mechanisms of the real sperm cell, we can treat our simple slender-filament model as defining a reduced dynamical phase space. The phase space is governed by the dimensionless sperm number 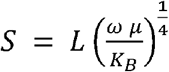 and the amplitude ratio 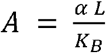, which control the emergence and structure of different beating patterns.

This suggests a practical assay protocol for positioning any observed flagellar system within the effective model’s phase space. First, the beating state of the flagellum is characterized experimentally using time-lapse imaging. Next, the dimensionless parameter *S* is estimated using measured values of flagellar length, beat frequency, fluid viscosity, and an assumed or measured bending stiffness. By systematically varying the external viscosity while keeping the other internal biochemical conditions fixed, one can identify the range of *S* values over which transitions into and out of three-dimensional beating occur. This defines a horizontal trajectory in the phase diagram. The position of the line that best matches the observed transition values of *S* allows one to assign the effective value of *A* to the flagellar system at hand. With the known or estimated value of the bending stiffness *K*_*B*_, we can also estimate the amplitude of the motor driving, α. This can then be used to estimate the maximum average force exerted by dynein heads in the flagellum considering that in a planar beat, the active moment on a cross section (per unit length), 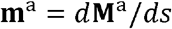 has a maximum magnitude of *αk*/*L* (see Methods), where *k* is the dimensionless wavenumber. At a cross section where the active moment is maximal, we expect that dynein heads on one half of the cross section (on 5 of the 9 microtubule doublets) are active (37), exerting forces longitudinally (perpendicular to the cross section). We consider that the longitudinal force per unit length at each microtubule, **f**^a^, is equal in magnitude on all 5 doublets, and that doublets are equidistant from the centre of the axoneme, so that only the angular positions of the doublets govern the contributions of these forces to the net active moment. This leads to an estimate of the magnitude of the net active moment at the cross section: |m^a^| ≈ 3 *r* |**f**^a^|, where *r* is the radius of the axoneme. Then the average force exerted by each dynein head can be estimated as *f*^*d*^ = |**f**^a^|/n = *A K*_*B*_ *k*/(3 *n r L*^2^) where *n* is the number of dynein heads per unit length on each microtubule. This approach assumes that motor activity is not modulated by external viscosity.

At present, there is no direct or non-destructive method to measure the average internal motor activity in the flagella of freely swimming sperm. This makes it difficult to quantify how motor regulation changes across species or physiological conditions using conventional approaches. The protocol described above to estimate *α* could be applied not only across different species but also across experimental conditions that affect internal regulation — such as ATP concentration, osmolarity, or drug treatments — thereby enabling a quantitative estimate of motor activity in sperm flagella.

We applied this idea to the sea-urchin sperm data of Ref. 6 and our samples of bull sperm. To determine the values of *S* corresponding to the viscosities in the experiments, for sea urchin sperm, we used a flagellar length of 42 µm, a beat frequency of 50 Hz, dimensionless beat wavenumber of 4π and a bending stiffness of 1 x 10^-21^ Nm^2^ (6, 38, 39). For bull sperm, we used a flagellar length of 66 µm, a beat frequency of 5 Hz, bending stiffness of 4x 10^-21^ Nm^2^ (40–42) and a dimensionless beat wavenumber of 2π from visual observation. The values of s corresponding to the viscosity values of 1 cP (planar), 1500 cP (helical) and 4000 cP (planar) in the sea urchin experiments are 3, 18 and 24, respectively, while values for the bull sperm in our experiments, *S* ≈ 2, 3, 4 and 6 for viscosities of 1 cP (quasiplanar), 5 cP (complex), 20 cP (planar) and 75 cP (planar), respectively. To ensure that beat transitions in our experiments are unaffected by medium viscoelasticity, we repeated the experiments with a PVP buffer restricted to the Newtonian regime and found that the viscosities at which beat transitions are observed remain the same (Supplementary Figure 3, Supplementary Videos 9 and 10). We then lined up the values of *S* and the observed motion states with the phase maps (Fig. 3A with *k* = 2π for bull sperm; Fig. 3B with *k* = 4π for sea urchin) to find the best values of *A* ≈ 5 for bull sperm and 30 for sea urchin sperm. This corresponds to α = 300 pN · μm and 710 pN · μm, respectively. Using an axoneme radius of *r* = 80 nm (43) and number of dynein heads (heavy chains) on a doublet per unit length, *n* ≈ 200 (44, 45), we obtain an average force per dynein head of 4.5 pN for sea urchin sperm and 0.6 pN for bull sperm.

The PSDs of the curvatures of the 2D-projections in the microscope images (Fig. 3 G, I, K) were found to be qualitatively comparable with the PSDs of the first time-dependent POD coefficient of curvature in the simulations (Fig. 3 H, J, L) with the parameter values estimated as discussed above.

### Emergence of 3D beat patterns consistent with mechanical instability

Our results above clearly demonstrate that even if the active bending moment in the axoneme always acts in the same plane in the local material reference frame associated with each cross section, the filament can develop torsion, and the beating can become non-planar. Further, in the regions of the phase map in Fig. 3A where the beating is 3D, the beating pattern quickly becomes non-planar from the straight-rod initial condition within a small (≲ 10) number of beat cycles (Supplementary Figure 4).

At low dimensionless frequencies, the emergent beat pattern closely follows the pattern of the internal driving. This behaviour can be understood by noting that when the dimensionless frequency *S* = 0, the form of the active moment resembles a time-independent “preferred curvature,” i.e., an equilibrium shape in which the stored elastic energy would be zero (see Methods). A force- and torque-free elastic filament displaced from such a state would naturally relax back to it. At small values of *S*, the filament experiences a slowly varying preferred curvature, which it can dynamically match as long as the forcing frequency is lower than the filament’s elastic relaxation rate. By construction, the forcing and elastic timescales become equal when *S* = 1.

Figure 4A shows the maximum difference between the actual curvature profile and the imposed dynamic preferred-curvature profile as a function of time for a range of dimensionless frequencies. These results are obtained using our nonlinear simulations. At low *S*, the error remains small, while at intermediate and high *S*, the filament does not match the preferred curvature. Figure 4B illustrates how this mismatch affects the filament’s shape over one beat cycle at a fixed amplitude *A* = 25 and wavenumber *k* = 4π. At high frequency (*S* = 18; top panel), the imposed waveform propagates too rapidly for the filament to respond elastically. As a result, the filament experiences the time-averaged effect of the fast-varying driving wave, which corresponds to a nearly straight configuration in the interior, with curvature localized near the ends due to boundary constraints. In contrast, at an intermediate frequency of *S*=9 (bottom panel), where the driving and response time scales are comparable, the filament develops a large-amplitude planar waveform.

**Fig. 4.**
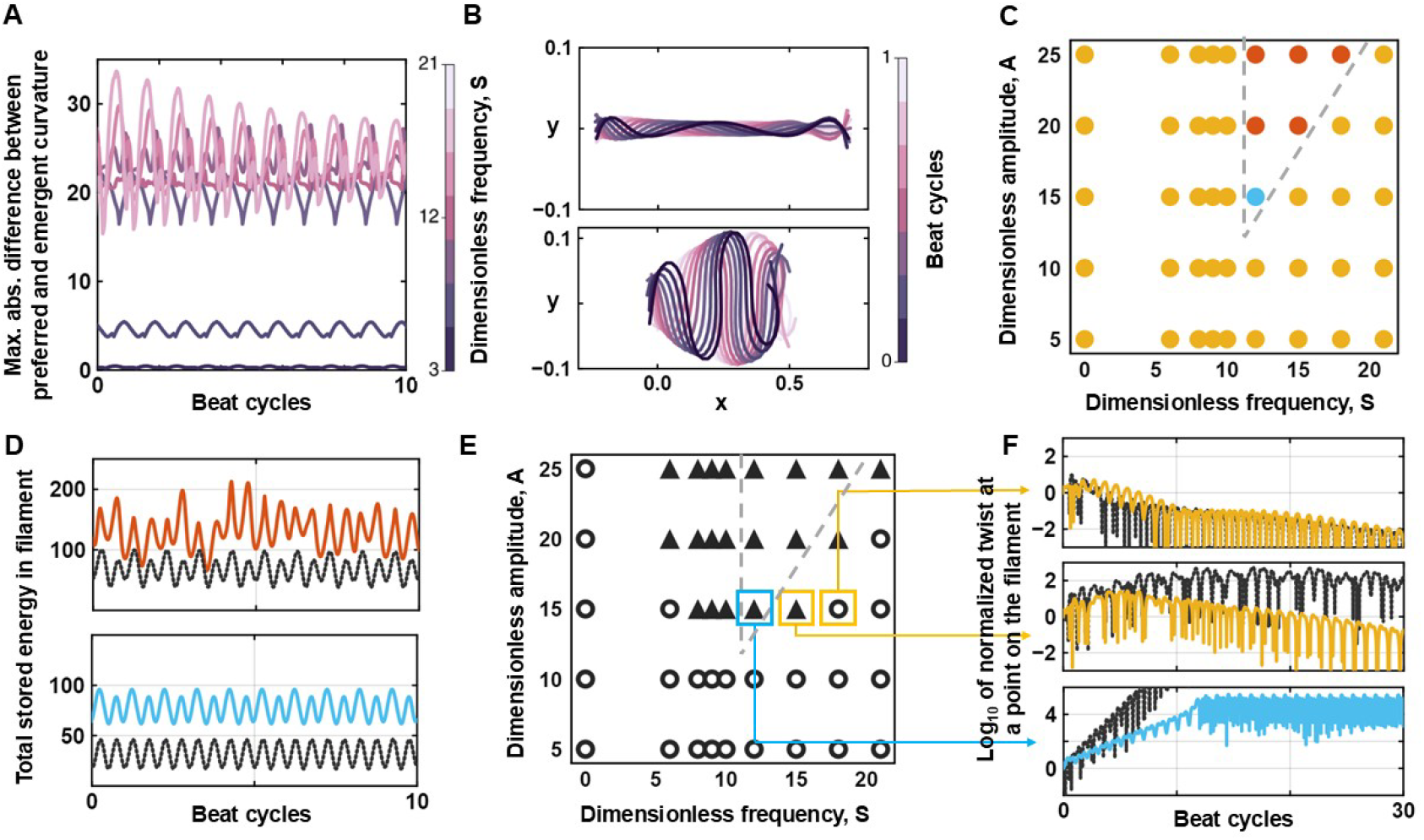
Elastohydrodynamic origins of the three-dimensional beats at k = 4π. (A) Maximum (along the arclength) of the absolute difference between the dynamic preferred curvature dictated by the internal driving and the filament’s emergent curvature at *A* = 20, *k* = 4π for different values of 5 (see colorbar). (B) Comparison of the shape of the filament over one beat cycle at a high dimensionless frequency, *S* = 18, *A* = 25, *k* = 4π (top) and intermediate dimensionless frequency, *S* = 9, *A* = 25, *k* = 4π (bottom). Each line represents a different time point (see colorbar). (C) Phase map showing the emergence of planar (yellow), helical (blue), and complex (red) beating when considering only local anisotropic hydrodynamic drag. (D) Comparison of the total energy over the filament as a function of time (in beat cycles) for the planar base state (black) and corresponding 3D beat (red or blue) at *S* = 12, *A* = 20, *k* = 4π (top) and *S* = 12, *A* = 15, *k* = 4π (bottom). Colors represent beat pattern types as in (C). (E) Phase map showing points where planar base states are linearly stable (circles) or unstable (triangles) to out-of-plane shape perturbations. Grey dashed lines enclose the region where 3D beating was observed in full nonlinear RFT simulations in (C). (F) Log_10_ of the twist at the filament midpoint (normalized with respect to the initial twist perturbation at t = 0) as a function of time for three points in the phase map in (E). Black dotted lines represent the linearized system results; colored lines correspond to nonlinear curvature-stress simulations (yellow represents planar beating, blue represents helical beating).

This leaves the question: why do beat patterns at intermediate dimensionless frequencies become non-planar? We first investigated whether the emergence of filament torsion requires the long-range hydrodynamic interactions generated by the flow around the filament. Our full simulations account for these interactions via Stokesian hydrodynamic-interaction tensors (37, 38). To isolate their influence, we repeated the simulations using the simpler Resistive Force Theory (RFT), which retains only the local anisotropic drag on each segment and neglects inter-segment flow coupling.

Interestingly, we found that three-dimensional (3D) beating still emerged in the RFT-based simulations (Fig. 4C), although the range of parameters over which it occurred was reduced compared to the full hydrodynamic model. The patterns observed in these simulations were predominantly complex, with only a single instance of helical beating. This indicates that the onset of non-planar motion does not require non-local hydrodynamic interactions, and may instead reflect an internally driven mechanical instability arising from the coupling between local drag anisotropy and filament elasticity.

At the same time, the prevalence of complex patterns in RFT — and the differences in pattern selection compared to simulations with full hydrodynamic interactions — suggest that long-range flow coupling may play a role in shaping the specific character of post-instability dynamics. In particular, the richness and diversity of beating patterns may depend on how fluid-mediated interactions distribute forces and correlations across the filament. Together, these results support the view that 3D transitions can arise from local mechanical effects alone, but the structure of the resulting waveform may be influenced by the nature of hydrodynamic coupling.

We then considered the possibility that the filament undergoes a buckling-like transition from planar to non-planar shapes due to emergent compressive forces. Buckling instabilities of this kind have been observed in many other systems involving actuated slender filaments, including microfilaments driven by ‘follower’ forces (39). To explore this, we used the RFT-based model and first restricted the system to two spatial dimensions, effectively suppressing out-of-plane deformations (see Methods). This 2D-constrained system produced stable planar beating solutions across the entire range of parameters tested, including those where 3D beating was observed in the full simulations.

This observation suggests that multiple steady-state solutions may exist for the same parameter values with planar and non-planar states coexisting, but only the latter is accessible when out-of-plane dynamics are permitted. In other words, the planar beating solution remains dynamically admissible in 2D but may become unstable to 3D perturbations in the full system. This is consistent with the idea that the transition to non-planar beating reflects a symmetry-breaking bifurcation, rather than a continuous deformation of the planar state. However, in contrast to classical buckling transitions, we find that the total elastic energy stored in the filament is higher in the 3D beating states compared to the corresponding planar beats at the same parameter values (Fig. 4D). In elastic rods under axial compression, buckling typically occurs when the deformed configuration reduces the system’s elastic energy while satisfying geometric and force constraints. Our system, by contrast, shows no such energetic advantage for the non-planar state. This suggests that the transition to 3D beating observed here is unlikely to be due to classical buckling, but instead reflects a non-equilibrium, symmetry-breaking mechanism.

Gadêlha et al. (2010) described a similar transition in flagellar beating as a buckling instability, driven by internally generated compressive forces. In their simulations, the onset of symmetry-breaking coincided with a drop in internal tension, which they interpreted as a signature of energy relief and hence buckling. Although elastic energy was not directly reported, their interpretation assumes that the deformed state is energetically favorable. In our case, by contrast, we observe a consistent increase in stored elastic energy in the 3D beating states. This key difference suggests that, although both systems exhibit spontaneous transitions to asymmetric waveforms, the underlying instability in our system does not conform to the classical energetic criteria for buckling.

Related studies of elastic filaments under follower forces or active driving have described rich dynamical transitions, including Hopf bifurcations and spontaneous three-dimensional motions (Ling et al., 2018). Other modeling frameworks have captured the coexistence of planar and non-planar solutions in elastohydrodynamic systems without necessarily invoking energy-based transitions (Gazzola et al., 2014). While these systems differ in detail, they support the broader view that active or resistively damped filaments can undergo symmetry-breaking instabilities not governed by energy minimization.

Taken together, our findings point to a form of driven mechanical instability that leads to non-planar dynamics without conferring energetic benefit. The spontaneous emergence of torsion, its dependence on forcing parameters, and the coexistence of planar and 3D solutions suggest a bifurcation-like non-equilibrium transition that is dynamical rather than energy-minimizing. While we do not attempt a full analytical classification here, the behavior we observe is not consistent with conventional buckling and may reflect a broader class of non-conservative instabilities in active slender filaments.

We analyzed the linear stability of planar beat patterns to small twist perturbations, which may either grow over successive beat cycles—leading to three-dimensional (3D) beating—or decay, resulting in a stable planar solution. To do this, we reformulated the governing equations in terms of curvature–stress variables and constructed a linearized system about the previously obtained planar base states (see Methods). The beat was represented as a combination of the base state and small leading-order perturbations. To test stability, we evolved the linearized system from initial conditions where all perturbations were set to zero for five beat cycles, followed by the introduction of a small twist perturbation. The subsequent evolution of twist was tracked for 30 beat cycles.

In addition to the linearized analysis, we also solved the full nonlinear curvature-stress equations for comparison. A useful feature of this formulation is that, when initialized with a perfectly planar configuration and no perturbation, the solution remains planar for all parameters, thus providing a controlled setting for perturbation analysis. Figure 4E shows the results of the linear stability tests: triangles indicate parameter values where twist perturbations grow, and circles indicate decay. The grey dashed lines enclose the region where 3D beating is observed in the earlier RFT simulations (Fig. 4C), allowing a comparison between linear predictions and fully nonlinear behavior.

We find that, at intermediate values of the dimensionless frequency *S*, many planar base states are linearly unstable to twist. However, only a subset of these unstable states— primarily those near the core of the unstable region—lead to sustained 3D beating in the fully nonlinear RFT simulations. Figure 4F compares the evolution of twist in the linearized system (black lines) and the nonlinear curvature–stress simulations (colored lines) at three representative points. In the top panel, twist decays in both cases, indicating stability. In the bottom panel, twist grows in both, indicating instability. In the middle panel, twist initially grows in the linear system, but then decays in the nonlinear system, suggesting that nonlinear effects stabilize the planar beat. This nonlinear suppression appears to be a common feature across many triangle-marked points outside the region bounded by the dashed lines in Fig. 4E. Together, these results indicate that the steady-state beating pattern at intermediate frequencies is determined by a competition between destabilizing linear modes and stabilizing nonlinear interactions.

### Beat patterns are strongly modified by flow near surfaces

Overall, the results in the sections above clearly show that the experimentally observed 2D-3D beating transitions in sperm are likely to be universally observable in any eukaryotic flagella, if the viscosities of the ambient fluid are such that the operating conditions span the phase boundaries in Fig. 3. The proximity of a no-slip surface also increases the hydrodynamic resistance encountered by a swimming cell. While our hydrodynamic model ignores lubrication interactions close to a wall, we can nevertheless explore how planar beats evolve as a swimmer approaches a surface.

Incorporating Stokesian wall hydrodynamic-interaction tensors into our 3D simulations, we found that a filament exhibiting steady planar beating in bulk (at *S* = 9, *A* = 12, *k* = 4π)) transitioned to a quasi-planar beat when it came within approximately 0.2 filament lengths (*L*) of an infinite plane wall. Figure 5A illustrates this transition: a filament initially placed parallel to the wall at a height of 1.5*L* was drawn toward the wall by hydrodynamic attraction (41). The beat pattern remained planar with a slight 1° tilt toward the wall (top inset), but as the filament accumulated at a distance of ∼0.18 *L* (∼8 μm for sea urchin sperm; ∼15 μm for bull or human sperm), the beat shifted to a quasi-planar mode (bottom inset). The power spectral density (Fig. 5B) reveals that this near-wall quasi-planar beat is qualitatively distinct from the quasi-planar beats seen in unbounded fluid (see Fig. 1N). Notably, the torsion kymograph (Fig. 5C) shows periodic sign reversals, corresponding to incomplete rolling alternations between clockwise and counterclockwise directions.

**Fig. 5.**
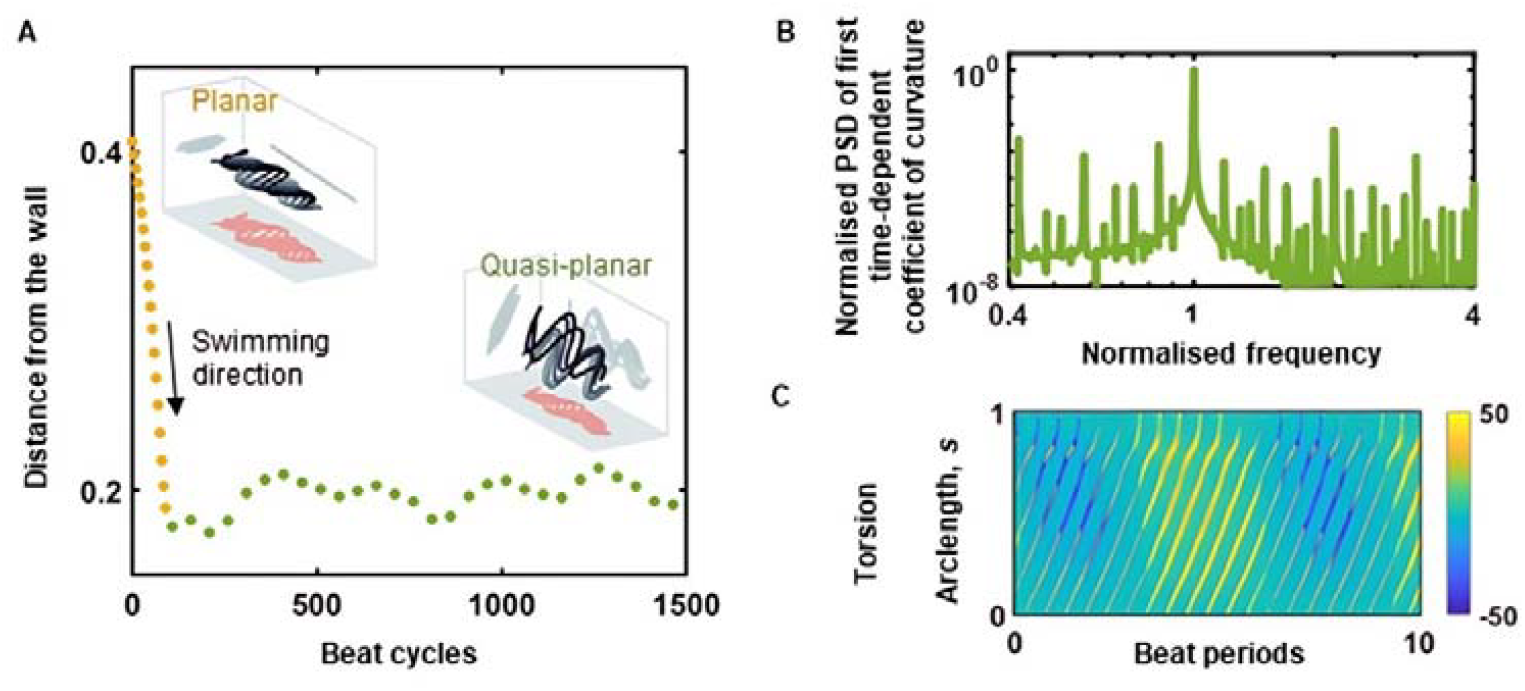
Transition to quasi-planar beating near a plane wall. (A) Head trajectory of an originally planar swimmer (*S* = 9, *A* = 12, *k* = 4π) near a wall, with a transition from a planar beat (top) to a quasi-planar beat (bottom). Yellow points denote time points with planar beating and green points indicate quasi-planar beating. (B) Normalised power spectral density of the first time-dependent coefficient of curvature in a quasi-planar beat. (C) Kymograph of the spatiotemporal evolution of centerline torsion in a quasi-planar beat.

Interestingly, this oscillatory roll-reversal behaviour resembles observations in sea urchin sperm swimming near a plane surface (6), where the cell rocks from side to side without completing full axial rolls. Importantly, no such changes were observed in simulations using RFT with resistance coefficients modified to account for wall proximity (42), indicating that the near-wall transition in beating pattern arises from the complex flow structure near the wall, which is not captured by local drag models.

## Discussion

The origin of non-planar beating patterns in eukaryotic cilia and flagella has been a long-standing mystery. It has been shown that spatiotemporal oscillations will spontaneously arise when motors act on elastic elements (4). This basic mechanism can be further augmented by the regulation of the motors (15, 43–45). Indeed, detailed experimental investigation of the states of the dynein motors has revealed that motors on one side of the doublet ring are inhibited at any time at each cross-section (46). These experiments also show that motors are switch-inhibited in an oscillatory manner as the bending wave propagates down the filament. These observations lend support to the idea that the wave of active bending moment generated collectively by the motors is internally planar. On the other hand, there is considerable experimental evidence that cilia and flagella exhibit both planar and non-planar beating patterns. To explain non-planar beating, more complex 3D regulation of motor activity has been suggested (6, 39, 47, 48). However, transitions between 2D and 3D beating are observed in a single species when the environmental conditions change. Explaining such transitions becomes challenging with mechanisms for 3D beat pattern generation that are purely based on motor regulation.

Our results instead show that 3D beating and 2D–3D beat transitions may have a simpler elastohydrodynamic origin that is generic to all eukaryotic species. This is consistent with the fact that the axonemal structure is highly conserved across eukaryotes, and that the underlying mechanisms giving rise to spontaneous bending-moment oscillations are also generic. The 3D wave patterns generated by this universal mechanism can capture the rolling, quasi-planar, helical, and more irregular beat patterns observed in experiments. Moreover, our results clearly show how the beating patterns depend on the length and stiffness of the ciliary or flagellar filament, the oscillatory characteristics of the active bending wave, and the hydrodynamic properties of the surrounding medium. We have shown that the variety of beating patterns arising from changes in ambient medium viscosity can be systematically predicted with a single model.

While the simulations and experimental comparisons in this study are specific to eukaryotic sperm flagella, the underlying elastohydrodynamic mechanism we identify arises from the conserved structure and universal motor-driven dynamics of the axoneme. It is therefore possible that similar transitions in beat patterns may also occur in motile cilia. However, given the shorter length scales, mechanical anchoring to cell surfaces, and distinct functional contexts of cilia, such extrapolations should be made cautiously and validated with system-specific models or experiments.

Our results appear consistent with the hypothesis that these transitions arise from an elastohydrodynamic instability. A full confirmation of this hypothesis requires a detailed Floquet stability analysis. Using the steady oscillatory planar beating states obtained with planar-constrained dynamics as the base states, we have already demonstrated here that small, arbitrary initial twist perturbations grow when integrated with the linearized equations about the base state. The Floquet analysis would reveal the nature of the Floquet multipliers in the unstable regime and thus enable a full bifurcation analysis of the identified eigenmodes analytically. Such analysis is expected to be the subject of future work.

This work aligns with recent studies that have emphasized the importance of mechanical instabilities in shaping the dynamics of cilia and flagella (11, 12, 14, 16, 17, 21, 23, 49–54). Gadêlha et al. (23) demonstrated that asymmetric beating in internally driven planar filaments can arise from buckling instabilities, without requiring morphological asymmetry or asymmetry in internal forces. Our results extend these findings to the emergence of three-dimensional asymmetric beating, also without invoking internal asymmetries. Furthermore, the beat patterns obtained here are robust to bending stiffness anisotropy on the order of the estimates provided by Lindemann and Lesich (35), which are based on a macroscopic reconstruction of the axoneme involving rigid adhesions, and may therefore be regarded as an upper bound in the absence of direct experimental measurements. This finding contrasts with results from a sliding-controlled model of the axoneme (55), where bending anisotropy led to a diversity of planar and three-dimensional beating patterns, whereas isotropic bending rigidities yielded only helical motion.

Our results also confirm earlier findings (including 16) that nonlinear effects become important at parameter values that induce high flagellar curvatures. Rode et al. (21) and Ling et al. (49) showed that mechanical instabilities can lead to transitions between planar and three-dimensional beating under different forms of internal driving. Both studies focused on how changes in internal driving parameters—such as the wavelength of the active bending moment in Rode et al. (21)—can produce such transitions. A purely internal mechanism for viscosity-related transitions would require some additional mechanosensory process to couple the internal forcing to the properties of the ambient medium. In contrast, our results show that viscosity-induced beat transitions in sperm could occur even in the absence of such complex internal regulation.

The emergence of three-dimensional waveforms from internally planar driving appears to be a robust outcome even when long-range non-local interactions are neglected and hydrodynamic drag is modelled using local anisotropic coefficients. This is relevant to experiments because experiments with sperm are almost always performed with sperm close to a surface. The proximity to a surface naturally screens long-range hydrodynamic interactions. The fact that we observe non-planar patterns even with locally anisotropic drag suggests that the same elastohydrodynamic instability mechanism may be responsible for the transitions observed experimentally with sperm swimming close to microscopic slide walls. Further, the differences between our simulations with and without long-range hydrodynamic interactions (Fig. 3B vs. Fig. 4C) show that these non-local interactions modify the waveforms that develop after the onset of instability. Our simulations of changes in beating patterns as filaments approach a wall are consistent with the fact that wall-induced screening progressively turns off the non-local modification. This is in line with experimental observations that beating patterns are altered by the presence of a wall (46). With sea urchin sperm (6), a transition from planar to quasi-planar beating – resembling a ‘flattened’ helical beat – was reported when moving away from the surface.

We have also demonstrated that the viscosity values at which beating transitions occur—from planar to quasi-planar and back—can be used to extract the dimensionless active moment amplitude, *A*, by matching experimental observations to simulation results. This value can then be related to the average force exerted by a dynein head in the axoneme. We obtained average force values of 4.5 pN and 0.6 pN for sea urchin sperm and bull sperm, respectively. These values are below the expected maximum value of 5 pN estimated from experiments (44). Our estimate for sea urchin sperm falls within the experimentally measured range of 2 – 5 pN per dynein head (44). While small differences in these values may be possible due to structural differences in dynein motors across species, the much smaller value we obtained for bull sperm is likely a result of modelling assumptions. Our assumptions of uniform flagellar thickness and bending stiffness particularly affect calculations for bull sperm, but not sea urchin sperm. The effect of including features such as the head remains to be fully understood (47, 48), but can be expected to lead to shifts in the transition boundaries, which would then affect the estimated dynein forcing.

The analysis above suggests that it is possible to exploit measurements of the viscosities at which 2D-3D transitions in beat patterns occur to extract quantitative estimates of internal forcing. Recently, a microfluidic device was used to systematically study the effect of varying viscosity on the flagellar waveforms of single sperm cells (42). Transitional viscosities obtained in such devices could be combined with independent measurements of the bending stiffness and compared with results from detailed simulations that account for features that have been neglected here (*e*.*g*. nonuniformity in filament radius due to the head and the flagellar taper, weak viscoelasticity induced by the methylcellulose viscosifier, *etc*.) to yield more accurate values of the dynein forcing. Such assays can study the effect of signalling molecules or ionic species on motor forcing, thus enabling the construction of detailed models of the biochemical regulation of the axoneme. Measurements of the response of dynein forcing to biochemical signals could also have diagnostic applications.

Interestingly, our experiments showed that despite the difference in wavenumber between sea urchin and bull sperm, the former exhibits planar beating at low viscosities, while the latter does so at high viscosities. The corresponding estimates of dynein force were also different between the two species, as were their lengths. It is possible that these parameters are tuned by evolution to enable planar beating in the viscosities typical of their natural environments. Because the dimensionless frequency *S* depends only weakly on viscosity (approximately as 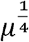), if the operating point of a given species lies far from the planar–helical transition boundary, its natural beating pattern may remain robust to large variations in the viscosity of the ambient fluid. On the other hand, if the operating point is close to the boundary, then small variations in filament length, morphology, or axonemal structure between individual sperm could lead to significantly different beating patterns, even within a single semen sample (7, 8).

Since the parameter *A* is directly proportional to the dynein forcing, biochemically mediated changes in motor force can shift the operating point vertically in phase space and thereby induce a transition in beating pattern and swimming trajectory. The implications of such changes for propulsion efficiency and metabolic cost remain to be explored.

## Methods

### Model for the flagellum

At any instant of time *t* the position and shape of the filament is characterized by the centerline position vector distribution, **r**(*s,t*), where *s* is the arclength coordinate along the centerline of the inextensible filament. The orientation of every cross-section is characterized by the triad of orthogonal unit vectors formed by the unit tangent vector,

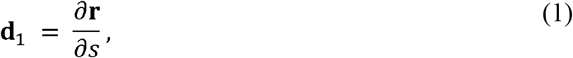

and the unit vectors, **d**_2_ and **d**_3_ that lie in the plane of the cross-section (Fig. 6). These material frame vectors are related to each other such that

**Fig. 6.**
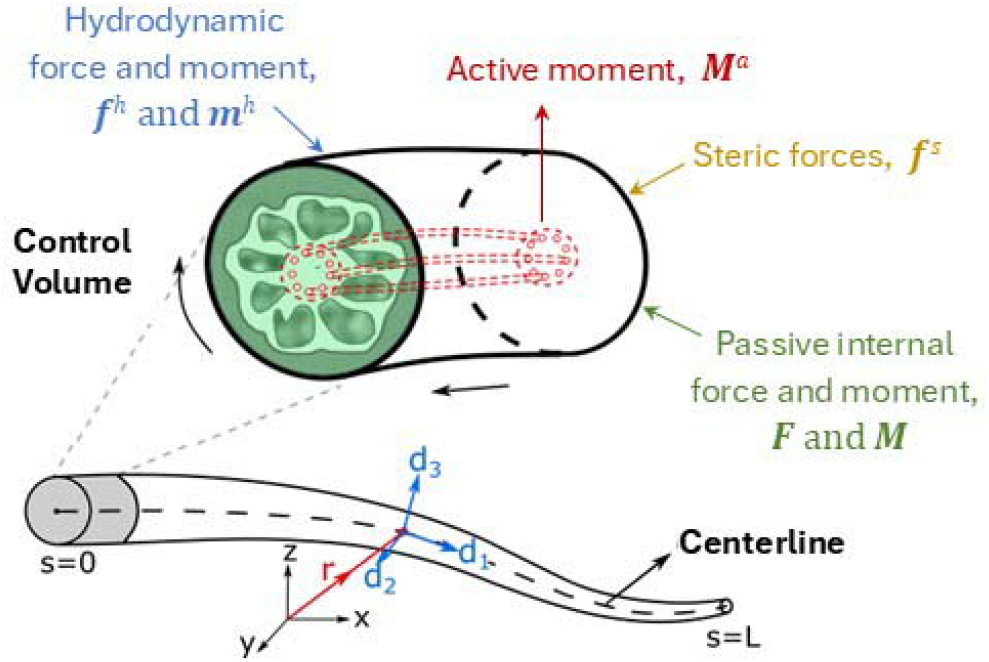
The internally-driven Kirchhoff-rod model.

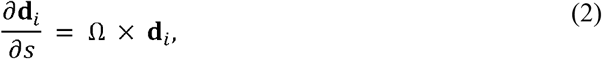

where the components of the Ω vector are the generalized curvatures that describe how quickly the material reference frame rotates as one travels down the filament, at any given instant of time (49).

The kinematics of the inextensible filament are given by:

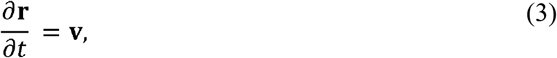

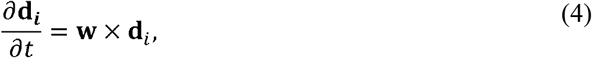

where **v** and **w** are the translational and rotational velocities of the rigid cross sections, respectively. Any differential control volume experiences external hydrodynamic forces and torques, and passive internal forces and moments due to elasticity that resist stretching, bending, and twisting. In the absence of any external forces, and when inertial effects are negligible, conservation of linear momentum yields (25, 50):

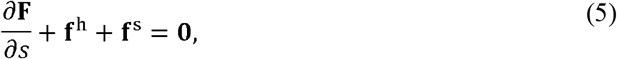

where **F** is the internal tension force exerted on a cross-section by the material on its aft side, and **f**^h^ and **f**^s^ are the hydrodynamic and steric forces per unit length. The steric excluded-volume force was implemented as in Schoeller *et al*. (51). Angular momentum conservation leads to (25, 50)

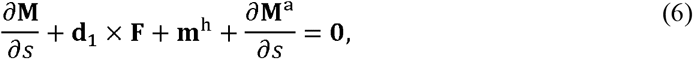

where **m**^h^ is the hydrodynamic moment per unit length. For a filament with no intrinsic curvature, the elastic bending moment at each cross-section is:

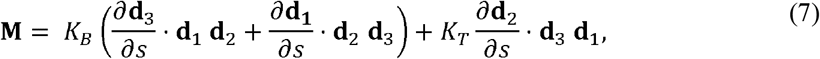

where *K*_*B*_ and *K*_*T*_ are bending and twisting stiffness coefficients, respectively. The effect of the axonemal driving is modeled as a travelling wave of active moment always acting about the binormal direction at every cross-section:

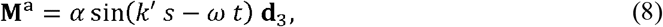

where *α,k*′ and *ω* are the amplitude, wavenumber, and frequency of the travelling wave. This driving moment is equivalent to imposing a travelling preferred-curvature wave in *Ω*_3_. The oscillating active moment at each cross-section is thus always exerted about the same axis in the material frame associated with that cross section. If the filament shape is always planar with respect to the global laboratory-attached reference frame, then the driving moment above will also be planar in the global reference frame. However, the kinematics and dynamics described by the set of model equations above do not constrain the filament to remain planar, and it is possible that the filament takes a non-planar shape in the global frame in response to the locally unidirectional active internal moment.

The relationship between the hydrodynamic force and moment and the linear and angular velocity at each cross section was expressed in terms of hydrodynamic mobility matrices as:

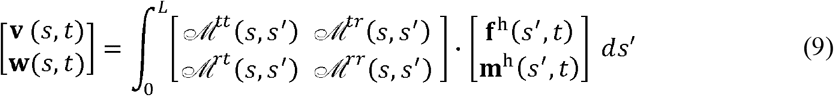

where ℳ^*tt*^, ℳ^*tr*^,ℳ^*rt*^ and ℳ^*rr*^ are mobility matrices that depend on the hydrodynamic model chosen. For filaments in an infinite viscous medium, long-range hydrodynamic interaction between different segments along the filament was modelled using the Rotne-Prager-Yamakawa mobility tensors (52, 53) the tensors provided by Swan and Brady (54) were used to model filaments near an infinite plane wall after ignoring lubrication effects. We also used Resistive Force Theory with the drag coefficients provided by Lighthill (55) for free space and coefficients by Katz *et al*. (56) near a wall. The forms of these mobility matrices are available in Supplementary Note 1.

Both ends of the filament are force and moment-free so that

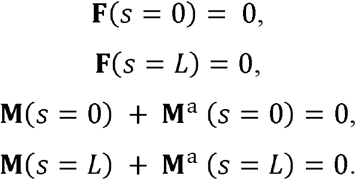

We used the numerical method developed by Schoeller *et al*. (51) to integrate the equations above forward in time until a steady oscillatory state was obtained. The material frame vectors were represented by unit quaternions to ensure that the vectors remain of unit magnitude at every time step, thus, satisfying the constraints that the filament is inextensible and cross-sections remain planar. An implicit second-order backward difference scheme was used to discretize the equations. Starting from a perfectly straight filament with **d**_*i*_ aligned with the unit vectors of the global reference frame, the quasi-Newton Broyden’s method was used to iteratively solve the nonlinear system of equations to calculate the internal tension **F** enforcing the inextensibility constraint, and **f**^h^ and **m**^h^ from the known shape at each time step. These were then used in Eqn. 9 to calculate the velocities, and then the translational and angular displacements at the end of that time step.

We used 1/*ω,L*, and *K*_*B*_*/L*^2^ as the time, length, and force scales, respectively. The dimensionless wavenumber of the forcing wave, *k* = *k*′ *L*, was set to either 2π or 4π to match experimental observations in bull and sea urchin. The swimming number, 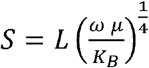, was varied between 2 and 21. The dimensionless active moment amplitude, 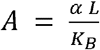, was varied between 5 and 30. The ratio of bending and twisting stiffnesses, *K*_*B*,_*/ K*_*T*_ was set to 1 and the dimensionless steric force parameter was set to a large value (10^2^). The filament was discretized into *N* = 50 segments of dimensionless length Δ*s* = 0.02 and radius α =0.01 units. This gives a radius-to-length ratio of 1/100, which approximately corresponds to that seen in various species of mammalian sperm, considering the radius of the thickest part of the flagellum (57).

### Analysis of beat patterns

We study shape changes using the Frenet curvature *κ*(*s,t*) and torsion *τ*(*s,t*). The curvature describes the local bending of the filament around any point, while the torsion describes the rate at which the centerline develops nonplanarity locally as it moves out of the local bending plane. These quantities remain invariant under any translation or rotation of the reference frame, and are defined as:

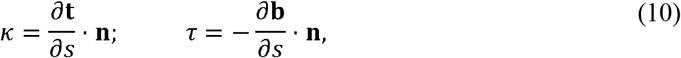

where **t** = **d**_1_ and **n** and **b** are the normal and binormal vectors, defined as:

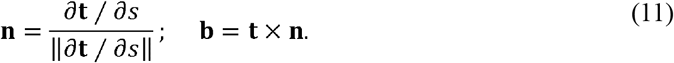

### Linear Stability Analysis

For generating oscillating planar base-states, only two of the material frame vectors change with time, and the shape of the filament was described by a single variable *θ*(*s,t*), such that for a filament beating in the x−y plane:

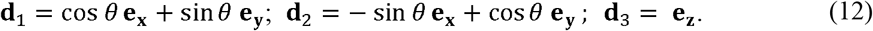

With this, the out-of-plane components of the translational velocities and forces are zero while angular velocities and torques have a component only in the **d**_**3**_ direction. The hydrodynamic force and moment are

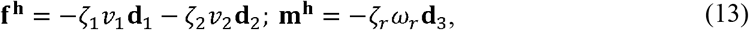

with the RFT drag coefficients (55):

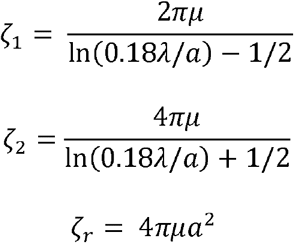

Since *Ω*_3_ is the only non-zero curvature component,

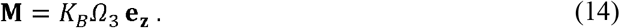

The system was numerically integrated in time using the numerical algorithm described in Schoeller *et al*. (51) to obtain planar base states at the required parameters.

To explore whether the planar base states predicted by the simulations above are linearly stable to perturbations in the shape, the model equations were recast in terms of the evolution of curvature as follows (see Supplementary Note 2):

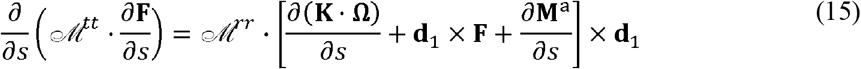

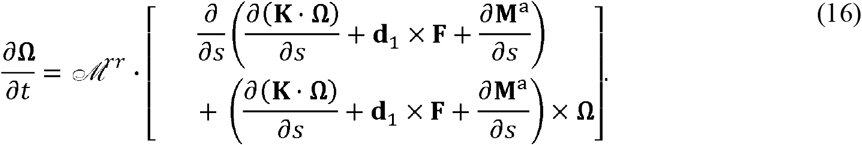

The curvature and internal stress were expanded as **Ω** = **Ω**^(0)^ + **Ω**^(1)^ + … and **F** = **F**^(0)^ + **F**^(1)^ + …, where **Ω**^(0)^ and **F**^(0)^ are the base-state curvature and internal tension predicted by the planar simulations, and **Ω**^(1)^ and **F**^(1)^represent leading order perturbations about the base state. By substituting these into Eqs. 15 and 16 and retaining terms only up to first order, we obtained the linearized scalar equations governing the evolution of the perturbation variables (see Supplementary Note 2). The perturbation variables were set to zero at the ends of the filament.

The base state and perturbation variables were first solved for simultaneously with all perturbation variables initialized to zero. After 5 beat cycles, a small twist perturbation was applied:

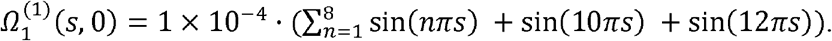

The PDE system was discretized automatically using the MethodOfLines.jl package in Julia v1.9.4 and integrated forward in time for 30 beat cycles using a Jacobian-free Newton Krylov algorithm available via the DifferentialEquations.jl package.

To study the effect of changing the initial condition, the initial twist perturbation defined above was multiplied by 0.8 and all other variables were kept the same:

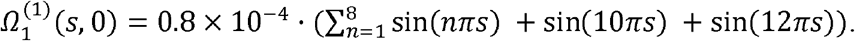

### Sperm Sample Preparation and Imaging

Bull semen was purchased from ABS Australia in 150*µL* straws and stored in liquid nitrogen prior to use. The semen samples were thawed in a 37°C water bath and diluted in a pre-warmed HEPES buffer (117 mM NaCl, 5.3 mM KCl, 1.8 mM CaCl_2_.2H_2_O, 0.8mM MgSO_4_, 1mM NaH_2_PO_4_, 5.5 mM D-Glucose, 0.03 mM Phenol Red, 4 mM NaHCO_3_, 21 mM HEPES, 0.33 mM Na Pyruvate, 21.4 mM Na Lactate, 0.3 mg/mL Bovine Serum Albumin and pH=7.4) supplemented with 0.1, 0.25 and 0.75% methylcellulose (M0512, Sigma-Aldrich) corresponding to a nominal viscosity of 1 cP, 5 cP and 75 cP, respectively. Diluted sperm samples were incubated at 37°C until imaging. To image bull sperm, 20*µL* of the prepared sperm sample were added to a chamber on a microscope slide, and then free-swimming sperm were imaged using high-speed high-resolution darkfield microscopy at 200 frames per second for 3 seconds. Imaging was performed using an AX-70 upright microscope (Olympus, Japan) equipped with an ORCA-Flash4.0 v2 sCMOS camera (Hamamatsu, Japan), a darkfield condenser (numerical aperture (NA)=0.9), and a U-DFA 18 mm internal diameter dark-field annulus with a UPlanAPO 10× magnification objective (NA= 0.4, Olympus, Japan). The sperm flagellar centreline was extracted using an automated image analysis algorithm as detailed in Nandagiri *et al*. (25).

## Data, Materials, and Software Availability

Microscopy data reported in this study are available within the article and its Supplementary Information files. Original code used to perform the simulations is publicly available at https://github.com/shibani-veeraragavan/Virtual-Flagellum.git. Any additional information required to reanalyze the data reported in this paper is available from the lead contact upon request.

## Acknowledgements

This work was supported by a Discovery Project Grant (DP190100343) from the Australian Research Council.

## Author contributions

S.V., R.P. and R.N. designed the research. S.V. and R.P. developed the mathematical model and analysis. S.V. implemented the code and performed the simulations. F.P. and R.N. designed the experiments. F.P. performed the experiments. S.V. analyzed data and wrote the manuscript. S.V., F.P., R.P. and R.N. edited the manuscript.

## Materials & Correspondence

Correspondence should be addressed to R.P. and R.N.

## References

1. J. L. Badano, N. Mitsuma, P. L. Beales, N. Katsanis, The Ciliopathies: An Emerging Class of Human Genetic Disorders. Annu Rev Genomics Hum Genet 7, 125–148 (2006).

2. A. Konno, M. Setou, K. Ikegami, “Ciliary and Flagellar Structure and Function—Their Regulations by Posttranslational Modifications of Axonemal Tubulin” in International Review of Cell and Molecular Biology, (Elsevier Inc., 2012), pp. 133–170.

3. P. Satir, T. Heuser, W. S. Sale, A Structural Basis for How Motile Cilia Beat. Bioscience 64, 1073–1083 (2014).

4. F. Jülicher, J. Prost, Spontaneous Oscillations of Collective Molecular Motors. Phys Rev Lett 78, 4510–4513 (1997).

5. R. Nosrati, A. Driouchi, C. M. Yip, D. Sinton, Two-dimensional slither swimming of sperm within a micrometre of a surface. Nat Commun 6, 8703 (2015).

6. D. M. Woolley, G. G. Vernon, A study of helical and planar waves on sea urchin sperm flagella, with a theory of how they are generated. Journal of Experimental Biology 204, 1333–1345 (2001).

7. T. Hyakutake, H. Suzuki, S. Yamamoto, Effect of viscosity on motion characteristics of bovine sperm. Journal of Aero Aqua Bio-mechanisms 4, 63–70 (2015).

8. M. Zaferani, F. Javi, A. Mokhtare, P. Li, A. Abbaspourrad, Rolling controls sperm navigation in response to the dynamic rheological properties of the environment. Elife 10 (2021).

9. P. Sartori, V. F. Geyer, A. Scholich, F. Jülicher, J. Howard, Dynamic curvature regulation accounts for the symmetric and asymmetric beats of Chlamydomonas flagella. Elife 5 (2016).

10. C. J. Brokaw, Computer Simulation of Flagellar Movement. Biophys J 12, 564–586 (1972).

11. D. Oriola, H. Gadêlha, J. Casademunt, Nonlinear amplitude dynamics in flagellar beating. R Soc Open Sci 4, 160698 (2017).

12. B. Chakrabarti, D. Saintillan, Spontaneous oscillations, beating patterns, and hydrodynamics of active microfilaments. Phys Rev Fluids 4, 43102 (2019).

13. I. H. Riedel□Kruse, A. Hilfinger, J. Howard, F. Jülicher, How molecular motors shape the flagellar beat. HFSP J 1, 192–208 (2007).

14. P. V. Bayly, K. S. Wilson, Equations of Interdoublet Separation during Flagella Motion Reveal Mechanisms of Wave Propagation and Instability. Biophys J 107, 1756–1772 (2014).

15. S. Camalet, F. Jülicher, Generic aspects of axonemal beating. New J Phys 2, 324 (2000).

16. C. Li, B. Chakrabarti, P. Castilla, A. Mahajan, D. Saintillan, Chemomechanical model of sperm locomotion reveals two modes of swimming. Phys Rev Fluids 8, 113102 (2023).

17. M. T. Gallagher, J. C. Kirkman-Brown, D. J. Smith, Axonemal regulation by curvature explains sperm flagellar waveform modulation. PNAS Nexus 2 (2023).

18. S. D. Olson, S. Lim, R. Cortez, Modeling the dynamics of an elastic rod with intrinsic curvature and twist using a regularized Stokes formulation. J Comput Phys 238, 169– 187 (2013).

19. L. Carichino, S. D. Olson, Emergent three-dimensional sperm motility: coupling calcium dynamics and preferred curvature in a Kirchhoff rod model. Math Med Biol 36, 439–469 (2019).

20. J. Simons, S. Olson, R. Cortez, L. Fauci, The dynamics of sperm detachment from epithelium in a coupled fluid-biochemical model of hyperactivated motility. J Theor Biol 354, 81–94 (2014).

21. S. Rode, J. Elgeti, G. Gompper, Sperm motility in modulated microchannels. New J Phys 21, 13016 (2019).

22. K. Ishimoto, E. A. Gaffney, An elastohydrodynamical simulation study of filament and spermatozoan swimming driven by internal couples. IMA J Appl Math 83, 655–679 (2018).

23. H. Gadêlha, E. A. Gaffney, D. J. Smith, J. C. Kirkman-Brown, Nonlinear instability in flagellar dynamics: a novel modulation mechanism in sperm migration? J R Soc Interface 7, 1689–1697 (2010).

24. E. H. Dill, Kirchhoff’s theory of rods. Arch Hist Exact Sci 44, 1–23 (1992).

25. A. Nandagiri, et al., Flagellar energetics from high-resolution imaging of beating patterns in tethered mouse sperm. Elife 10 (2021).

26. A. L. Hall-McNair, T. D. Montenegro-Johnson, H. Gadêlha, D. J. Smith, M. T. Gallagher, Efficient implementation of elastohydrodynamics via integral operators. Phys Rev Fluids 4, 113101 (2019).

27. S. Werner, J. C. Rink, I. H. Riedel-Kruse, B. M. Friedrich, Shape Mode Analysis Exposes Movement Patterns in Biology: Flagella and Flatworms as Case Studies. PLoS One 9, e113083 (2014).

28. F. Ling, H. Guo, E. Kanso, Instability-driven oscillations of elastic microfilaments. J R Soc Interface 15, 20180594 (2018).

29. K. Ishimoto, H. Gadêlha, E. A. Gaffney, D. J. Smith, J. Kirkman-Brown, Human sperm swimming in a high viscosity mucus analogue. J Theor Biol 446, 1–10 (2018).

30. D. J. Smith, E. A. Gaffney, H. Gadêlha, N. Kapur, J. C. Kirkman□Brown, Bend propagation in the flagella of migrating human sperm, and its modulation by viscosity. Cell Motil 66, 220–236 (2009).

31. A. Bukatin, I. Kukhtevich, N. Stoop, J. Dunkel, V. Kantsler, Bimodal rheotactic behavior reflects flagellar beat asymmetry in human sperm cells. Proceedings of the National Academy of Sciences 112, 15904–15909 (2015).

32. B. M. Friedrich, I. H. Riedel-Kruse, J. Howard, F. Jülicher, High-precision tracking of sperm swimming fine structure provides strong test of resistive force theory. Journal of Experimental Biology 213, 1226–1234 (2010).

33. A. Gong, et al., Reconstruction of the three-dimensional beat pattern underlying swimming behaviors of sperm. The European Physical Journal E 44, 87 (2021).

34. G. G. Vernon, D. M. Woolley, Three-dimensional motion of avian spermatozoa. Cell Motil Cytoskeleton 42, 149–161 (1999).

35. J. C. Kirkman-Brown, D. J. Smith, Sperm motility: is viscosity fundamental to progress? Mol Hum Reprod 17, 539–544 (2011).

36. C. B. Lindemann, K. A. Lesich, Functional anatomy of the mammalian sperm flagellum. Cytoskeleton 73, 652–669 (2016).

37. J. Lin, D. Nicastro, Asymmetric distribution and spatial switching of dynein activity generates ciliary motility. Science (1979) 360 (2018).

38. M. Okuno, Y. Hiramoto, Direct Measurements of the Stiffness of Echinoderm Sperm Flagella. Journal of Experimental Biology 79, 235–243 (1979).

39. J. Gray, G. J. Hancock, The Propulsion of Sea-Urchin Spermatozoa. Journal of Experimental Biology 32, 802–814 (1955).

40. N.-H. Gu, W.-L. Zhao, G.-S. Wang, F. Sun, Comparative analysis of mammalian sperm ultrastructure reveals relationships between sperm morphology, mitochondrial functions and motility. Reproductive Biology and Endocrinology 17, 66 (2019).

41. C. B. Lindemann, W. G. Rudd, R. Rikmenspoel, The Stiffness of the Flagella of Impaled Bull Sperm. Biophys J 13, 437–448 (1973).

42. F. Yazdan Parast, et al., Viscous Loading Regulates the Flagellar Energetics of Human and Bull Sperm. Small Methods (2023). 10.1002/smtd.202300928.

43. H. M. Sakakibara, Y. Kunioka, T. Yamada, S. Kamimura, Diameter Oscillation of Axonemes in Sea-Urchin Sperm Flagella. Biophys J 86, 346–352 (2004).

44. C. B. Lindemann, Structural-Functional Relationships of the Dynein, Spokes, and Central-Pair Projections Predicted from an Analysis of the Forces Acting within a Flagellum. Biophys J 84, 4115–4126 (2003).

45. K. Ishibashi, H. Sakakibara, K. Oiwa, Force-Generating Mechanism of Axonemal Dynein in Solo and Ensemble. Int J Mol Sci 21, 2843 (2020).

46. E. A. Gaffney, K. Ishimoto, B. J. Walker, Modelling Motility: The Mathematics of Spermatozoa. Front Cell Dev Biol 9 (2021).

47. K. Ishimoto, E. A. Gaffney, A study of spermatozoan swimming stability near a surface. J Theor Biol 360, 187–199 (2014).

48. K. Ishimoto, Hydrodynamic evolution of sperm swimming: Optimal flagella by a genetic algorithm. J Theor Biol 399, 166–174 (2016).

49. T. R. Powers, Dynamics of filaments and membranes in a viscous fluid. Rev Mod Phys 82, 1607–1631 (2010).

50. M. Hines, J. J. Blum, Bend propagation in flagella. I. Derivation of equations of motion and their simulation. Biophys J 23, 41–57 (1978).

51. S. F. Schoeller, A. K. Townsend, T. A. Westwood, E. E. Keaveny, Methods for suspensions of passive and active filaments. J Comput Phys 424, 109846 (2021).

52. E. Wajnryb, K. A. Mizerski, P. J. Zuk, P. Szymczak, Generalization of the Rotne– Prager–Yamakawa mobility and shear disturbance tensors. J Fluid Mech 731, R3 (2013).

53. L. Durlofsky, J. F. Brady, G. Bossis, Dynamic simulation of hydrodynamically interacting particles. J Fluid Mech 180, 21 (1987).

54. J. W. Swan, J. F. Brady, Simulation of hydrodynamically interacting particles near a no-slip boundary. Physics of Fluids 19, 113306 (2007).

55. J. Lighthill, Flagellar Hydrodynamics. SIAM Review 18, 161–230 (1976).

56. D. F. Katz, J. R. Blake, S. L. Paveri-Fontana, On the movement of slender bodies near plane boundaries at low Reynolds number. J Fluid Mech 72, 529 (1975).

57. N.-H. Gu, W.-L. Zhao, G.-S. Wang, F. Sun, Comparative analysis of mammalian sperm ultrastructure reveals relationships between sperm morphology, mitochondrial functions and motility. Reproductive Biology and Endocrinology 17, 66 (2019).

