## Supplementary Text for "Elastohydrodynamic mechanisms govern beat pattern transitions in eukaryotic flagella"

**The Supplementary Information for this manuscript includes:**

Supplementary Notes 1 and 2

Supplementary Figures 1 – 4

Description of Supplementary Movies S1 to S7 (separate files)

### Supplementary Figure 1

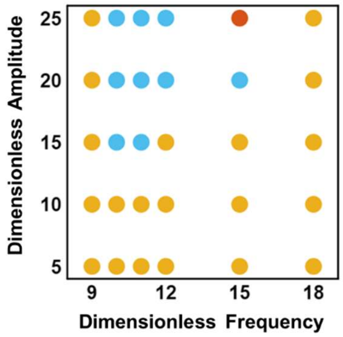


**Supplementary Figure 1 | Effect of bending stiffness anisotropy on beating patterns.** Phase map showing the emergence of planar (yellow), helical (blue) and complex (red) beating at $k = 4\pi$ when considering anisotropy in bending stiffnesses such that $K_{B,3}/K_{B,2} = 2.6$, where $K_{B,3}$ and $K_{B,2}$ are the bending stiffnesses along $\boldsymbol{d}_3$ and $\boldsymbol{d}_2$, respectively.

### Supplementary Figure 2

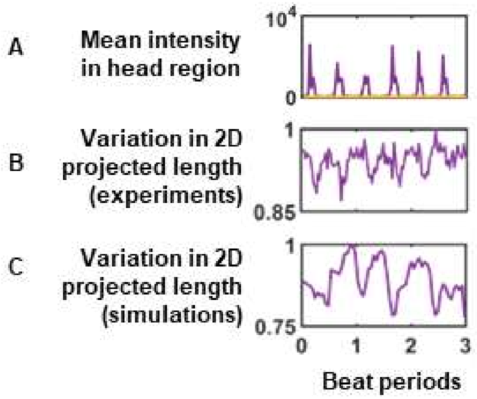


**Supplementary Figure 2 | Beat patterns of bull sperm at different viscosities.** (A) Mean light intensity over the head region of a quasiplanar (purple line) and planar (yellow line) bull sperm over 3 beat periods, obtained from microscope videos. (B) Variation in the 2D projected length of the quasiplanar bull sperm, based on the plane of view shown in **Error! Reference source not found.**G. (C) Variation in the 2D projected length of a simulated quasiplanar sperm at $S = 5, A = 5, k = 2\pi$.

### Supplementary Figure 3

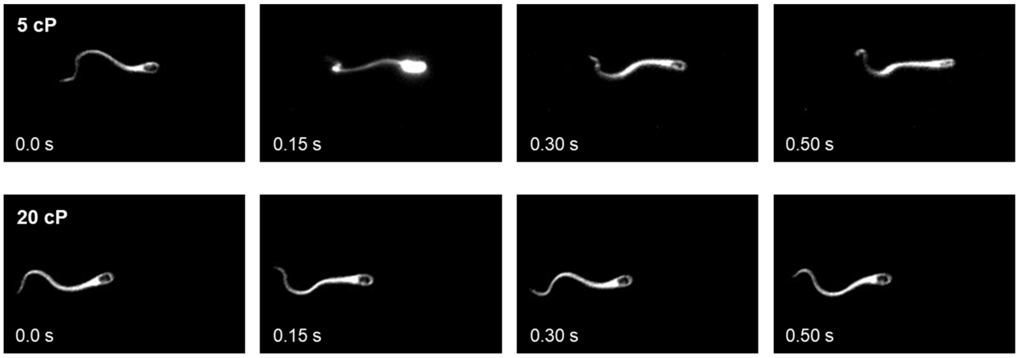


**Supplementary Figure 3 | Beating patterns of bull sperm in Newtonian PVP media.** Snapshots of the beating pattern at four timepoints in (top panel) a complex beat observed at 5 cP and (bottom panel) planar beat at 20 cP. Beat three-dimensionality is indicated by a change in light intensity at the head.

### Supplementary Figure 4

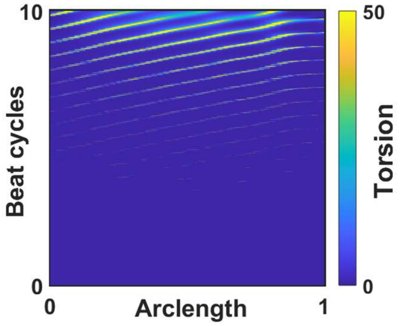


**Supplementary Figure 4 | Gradual emergence of torsion in a non-planar beat.** Kymograph of centerline torsion over 10 beat cycles for a helical swimmer at $S = 10, A = 15, k = 4\pi$.

#### Supplementary Note 1

The hydrodynamic mobility matrices in free space are written as follows (50, 51) and arranged into the form of a grand mobility matrix:

$$\mathcal{M}_{mn}^{tt} = \begin{cases} \dfrac{1}{8\pi\mu x}\left[\left(1+\dfrac{2a^2}{3x^2}\right)\mathbf{I}+\left(1-\dfrac{2a^2}{x^2}\right)\hat{\mathbf{x}}\hat{\mathbf{x}}\right] & ,m \neq n \\ \dfrac{1}{6\pi\mu a}\mathbf{I} & ,m = n \end{cases} \tag{S1}$$

$$\mathcal{M}_{mn}^{tr} = \mathcal{M}_{mn}^{rt\,\dagger} = \begin{cases} \dfrac{1}{8\pi\mu x^2}\ \boldsymbol{\varepsilon}\cdot\hat{\mathbf{x}} & ,m \neq n \\ \mathbf{0} & ,m = n \end{cases} \tag{S2}$$

$$\mathcal{M}_{mn}^{rr} = \begin{cases} \dfrac{1}{16\pi\mu x^3}\ (\mathbf{3}\,\hat{\mathbf{x}}\hat{\mathbf{x}} - \mathbf{I}) & ,m \neq n \\ \dfrac{1}{8\pi\mu a^3}\ \mathbf{I} & ,m = n \end{cases} \tag{S3}$$

where $\mu$ is fluid viscosity, $\mathbf{x} = \mathbf{r}_m - \mathbf{r}_n$ for two segments *m* and *n* with position vectors $\mathbf{r}_m$ and $\mathbf{r}_n$, $x = |\mathbf{x}|$, $\hat{\mathbf{x}} = \mathbf{x}/x$, $\boldsymbol{\varepsilon}$ is the three-dimensional Levi-Civita symbol and $\mathbf{I}$ is the identity tensor. The mobility matrices for motion near a wall have a more complicated form and are available in Swan and Brady (52). We do not include sphere-sphere or sphere-wall lubrication interactions. When using Resistive Force Theory with Lighthill's free-space coefficients (53), the mobility matrices are written as:

$$\mathcal{M}_{mn}^{tt} = \frac{\delta_{mn}}{\Delta s}\left(\frac{\ln(0.18\lambda/a)-1/2}{2\pi\mu}\mathbf{d}_1\mathbf{d}_1 + \frac{\ln(0.18\lambda/a)+1/2}{4\pi\mu}(\mathbf{d}_2\mathbf{d}_2+\mathbf{d}_3\mathbf{d}_3)\right), \tag{S4}$$

$$\mathcal{M}_{mn}^{tr} = \mathcal{M}_{mn}^{rt\,\dagger} = \mathbf{0}\,, \tag{S5}$$

$$\mathcal{M}_{mn}^{rr} = \frac{\delta_{mn}}{8\pi\mu a^3}\ \mathbf{I}\,, \tag{S6}$$

where $\delta_{mn}$ is the Kronecker delta, $\Delta s$ is the length of the segment and $\lambda$ is the beat wavelength. Near a wall we use the coefficients suggested by Katz *et al.* (54) and write the matrices as follows, where *h* is the distance between the segment and the wall:

$$\mathcal{M}_{mn}^{tt} = \frac{\delta_{mn}}{\Delta s}\left(\frac{ln(2h/a)}{2\pi\mu}\ \boldsymbol{d}_1\boldsymbol{d}_1 + \frac{ln(2h/a)}{4\pi\mu}\ \boldsymbol{d}_2\boldsymbol{d}_2 + \frac{ln(2h/a)-1}{4\pi\mu}\ \boldsymbol{d}_3\boldsymbol{d}_3\right), \tag{S7}$$

$$\mathcal{M}_{mn}^{tr} = \mathcal{M}_{mn}^{rt\,\dagger} = \mathbf{0}\,, \tag{S8}$$

$$\mathcal{M}_{mn}^{rr} = \frac{\delta_{mn}}{8\pi\mu a^3}\ \boldsymbol{I}\,. \tag{S9}$$

#### Supplementary Note 2

To derive the linearized equations describing the evolution of perturbations in curvature, we began by casting our equations into the curvature-stress form (Eqs. 15-16 in the Main Text). This may be derived as described below.

When using local drag theory, ignoring steric forces, the linear and angular velocity can be expressed as follows:

$$\mathbf{v} = \mathcal{M}^{\boldsymbol{tt}} \cdot \mathbf{f}^h \tag{S10}$$

$$\mathbf{w} = \mathcal{M}^{\boldsymbol{rr}} \cdot \mathbf{m}^h \tag{S11}$$

where $\mathcal{M}^{\boldsymbol{tt}}$ and $\mathcal{M}^{\boldsymbol{rr}}$ are the hydrodynamic mobility matrices defined in Eqs. S4-S6. Substituting the linear and angular momentum balances (Eqs. 5-6 in the Main Text) into these equations, we obtain:

$$\mathbf{v} = -\mathcal{M}^{\boldsymbol{tt}} \cdot \frac{\partial \mathbf{F}}{\partial s}, \tag{S12}$$

$$\mathbf{w} = -\mathcal{M}^{rr} \cdot \left(\frac{\partial \mathbf{M}}{\partial s} + \mathbf{d}_1 \times \mathbf{F} + \frac{\partial \mathbf{M}^{\mathrm{a}}}{\partial s}\right). \tag{S13}$$

Now, from the definition of the tangent (Eq. 1 in the Main Text) we have that

$$\frac{\partial}{\partial s}\left(\frac{\partial \mathbf{r}}{\partial t}\right) = \frac{\partial}{\partial t}\left(\frac{\partial \mathbf{r}}{\partial s}\right) = \frac{\partial \mathbf{d}_1}{\partial t}, \tag{S14}$$

and from Eq. 4 in the Main Text,

$$\frac{\partial \mathbf{d}_1}{\partial t} = \mathbf{w} \times \mathbf{d}_1. \tag{S15}$$

Substituting Eq. S12 and S13 into Eq. S15, we obtain Eq. 15 from the Main Text:

$$\frac{\partial}{\partial s}\left(\mathcal{M}^{tt} \cdot \frac{\partial \mathbf{F}}{\partial s}\right) = \mathcal{M}^{rr} \cdot \left[\frac{\partial(\mathbf{K} \cdot \boldsymbol{\Omega})}{\partial s} + \mathbf{d}_1 \times \mathbf{F} + \frac{\partial \mathbf{M}^{\mathrm{a}}}{\partial s}\right] \times \mathbf{d}_1. \tag{S16}$$

To obtain Eq. 16, we substitute Eq. S13 into the compatibility relation (47)

$$\frac{\partial \boldsymbol{\Omega}}{\partial t} = \frac{\partial \mathbf{w}}{\partial s} + \mathbf{w} \times \boldsymbol{\Omega} \tag{S17}$$

to obtain

$$\frac{\partial \Omega}{\partial t} = \mathcal{M}^{rr} \cdot \left[\frac{\partial}{\partial s}\left(\frac{\partial(\mathbf{K} \cdot \Omega)}{\partial s} + \mathbf{d}_1 \times \mathbf{F} + \frac{\partial \mathbf{M}^{\mathrm{a}}}{\partial s}\right) + \left(\frac{\partial(\mathbf{K} \cdot \Omega)}{\partial s} + \mathbf{d}_1 \times \mathbf{F} + \frac{\partial \mathbf{M}^{\mathrm{a}}}{\partial s}\right) \times \Omega\right]. \tag{S18}$$

We note that analogous expressions have been derived previously (15). The curvature and internal stress were then expanded as $\mathbf{\Omega} = \mathbf{\Omega}^{(0)} + \mathbf{\Omega}^{(1)} + \cdots$ and $\mathbf{F} = \mathbf{F}^{(0)} + \mathbf{F}^{(1)} + \cdots$, where $\mathbf{\Omega}^{(0)}$ and $\mathbf{F}^{(0)}$ are the base-state curvature and internal tension predicted by the planar simulations, and $\mathbf{\Omega}^{(1)}$ and $\mathbf{F}^{(1)}$ represent leading order perturbations about the base state. By substituting these into Eqs. S16 and S18 and retaining terms only up to first order, we obtained:

$$F_1^{1''} - \frac{\zeta_1}{\zeta_3}\left(2F_1^o \Omega_3^o \Omega_3^1 + F_1^1 \Omega_3^{o\,2}\right) - \left(1 + \frac{\zeta_1}{\zeta_2}\right)\left(\Omega_3^o F_2^{1'} + \Omega_3^1 F_2^{o'}\right) - F_2^o \Omega_3^{1'} - F_2^1 \Omega_3^{o'} = 0,$$

$$F_2^{1''} - \frac{\zeta_2}{\zeta_1}\left(2F_2^o \Omega_3^o \Omega_3^1 + F_2^1 \Omega_3^{o\,2}\right) + \left(1 + \frac{\zeta_3}{\zeta_1}\right)\left(\Omega_3^o F_1^{1'} + \Omega_3^1 F_1^{o'}\right) + F_1^o \Omega_3^{1'} + F_1^1 \Omega_3^{o'} -$$
$$\frac{\zeta_2}{\zeta_r}\left(K_B \Omega_3^{1'} + F_2^1 + M^{a'}\right) = 0,$$

$$F_2^{1''} - \left(1 + \frac{\zeta_2}{\zeta_1}\right)\Omega_2^1 F_1^{o'} - F_1^o \Omega_2^{1'} + F_1^o \Omega_3^o \Omega_1^1 + \frac{\zeta_2}{\zeta_1} F_2^o \Omega_3^o \Omega_2^1 + 2\Omega_1^1 F_2^{o'} + F_2^o \Omega_1^{1'} + \frac{\zeta_3}{\zeta_r}\left(K_B \Omega_2^{1'} -\right.$$
$$\left. F_3^1 - \Omega_1^1 M^{a'}\right) = 0,$$

$$-\zeta_r \dot{\Omega}_1^1 + K_T \Omega_1^{1''} + M^{a'} \Omega_2^1 + M_0^a \Omega_2^{1'} + K_B \Omega_2^1 \Omega_3^{o'} + F_2^o \Omega_2^1 + \Omega_2^1 M^{a'} - K_B \Omega_3^o \Omega_2^{1'} +$$
$$M^a \Omega_3^o \Omega_1^1 + \Omega_3^o F_3^1 = 0,$$

$$-\zeta_r \dot{\Omega}_2^1 + K_B \Omega_2^{1''} - M^{a'} \Omega_1^1 - M^a \Omega_1^{1'} - F_3^{1'} - K_B \Omega_1^1 \Omega_3^{o'} - F_2^o \Omega_1^1 - \Omega_1^1 M^{a'} + K_T \Omega_3^o \Omega_1^{1'} +$$
$$M^a \Omega_3^o \Omega_2^1 = 0,$$

$$-\zeta_r \dot{\Omega}_3^1 + K_B \Omega_3^{1''} + M^{a''} + F_2^{1'} = 0,$$

with $M^a = \alpha \cos(ks - \omega t)$, the primes denoting differentiation with respect to $s$, and the dots denoting differentiation with respect to $t$. Here $\zeta_1$, $\zeta_2$, $\zeta_3$ and $\zeta_r$ denote the RFT resistance coefficients such that:

$$\mathcal{M}^{tt} = \frac{1}{\zeta_1}\,\mathbf{d}_1\mathbf{d}_1 + \frac{1}{\zeta_2}\,\mathbf{d}_2\mathbf{d}_2 + \frac{1}{\zeta_3}\,\mathbf{d}_3\mathbf{d}_3\,,$$

$$\mathcal{M}^{rr} = \frac{1}{\zeta_r}\,\mathbf{I}\,.$$

#### Description of Supplementary Movies

**Movie S1.**

Planar beating pattern obtained in our simulations at $S = 7, A = 5, k = 2\pi$. Colors represent lighting such that a point higher in the *z* direction has a lighter color.

**Movie S2.**

Quasi-planar beating pattern obtained in our simulations at $S = 5, A = 5, k = 2\pi$. Colors represent lighting such that a point higher in the *z* direction has a lighter color.

**Movie S3.**

Helical beating pattern obtained in our simulations at $S = 12, A = 20, k = 4\pi$. Colors represent lighting such that a point higher in the *z* direction has a lighter color.

**Movie S4.**

Complex beating pattern obtained in our simulations at $S = 15, A = 20, k = 4\pi$. Colors represent lighting such that a point higher in the *z* direction has a lighter color.

**Movie S5.**

Quasi-planar beating pattern observed in our experiments of bull sperm at 1 cP.

**Movie S6.**

Complex beating pattern observed in our experiments of bull sperm at 5 cP.

**Movie S7.**

Planar beating pattern observed in our experiments of bull sperm at 20 cP.
